# C. perfringens enterotoxin-claudin pore complex: Models for structure, mechanism of pore assembly and cation permeability

**DOI:** 10.1101/2024.08.02.605804

**Authors:** Santhosh Kumar Nagarajan, Joy Weber, Daniel Roderer, Jörg Piontek

## Abstract

The pore-forming *Clostridium perfringens* enterotoxin (CPE), a common cause of foodborne diseases, facilitates Ca^2+^ influx in enterocytes, leading to cell damage. Upon binding to certain claudins (e.g. claudin-4), CPE forms oligomeric pores in the cell membrane. While the CPE-claudin interaction mechanism is well known, the structure and assembly of the pore complex remain elusive. Here, using AlphaFold2 complex prediction, structure alignment, and molecular dynamics simulations, we generated models of pre-pore and pore states of the CPE-claudin-4 complex sequentially addressing CPE-claudin, CPE-CPE, and claudin-claudin interactions, along with CPE conformational changes. The CPE pore is a hexameric variation of the typically heptameric pore stem and cap architecture of aerolysin-like β-barrel pore-forming toxins (β-PFT). The pore is lined with three hexa-glutamate rings, which differs from other β-PFTs and confers CPE-specific cation selectivity of the pore. Additionally, the pore center is indicated to be anchored by a dodecameric claudin ring formed by a cis-interaction variant of an interface found in claudin-based tight junction strands. Mutation of an interfacial residue inhibited CPE-mediated cell damage *in vitro*. We propose that this claudin ring constitutes an anchor for a twisting mechanism that drives extension and membrane insertion of the CPE β-hairpins. Our pore model agrees with previous key experimental data providing insights into the structural mechanisms of CPE-mediated cytotoxic cation influx.

## Introduction

*Clostridium perfringens*, an anaerobic, Gram-positive spore-forming bacterium, is known to cause a variety of illnesses worldwide, such as gas gangrene, wound infections, and various diseases affecting the gastrointestinal system of both humans and animals.^1^ Its toxicity derives from a series of at least 16 toxins produced by different strains.^2^

Infection with *C. perfringens* type F (formerly type A) is responsible for one of the most common foodborne diseases worldwide caused by release of the *C. perfringens* enterotoxin (CPE).^3, 4^ CPE has been shown to cause characteristic gastrointestinal symptoms.^5^ These are typically short-term but can result in serious complications, such as lethal necrotizing colitis.^6^ In addition, CPE was identified to play a role in non-foodborne diseases like antibiotic-associated diarrhea^7^, similar to C*lostridioles difficile.*^8^

After bacteria ingestion, usually through consumption of contaminated food, the sporulation process takes place in the intestines and ends with lysis of the mother cell. Thereby, previously over-expressed enterotoxin is released.^2, 9^ CPE binds to intestinal enterocytes through receptors, which are certain claudins (CLDNX), such as CLDN3 and -4.^10, 11^ Claudins, a ≥27-member family of ∼23-34 kDa proteins with four transmembrane helices (TM) and two extracellular segments (ECS), form the backbone of tight junctions, and regulate the paracellular permeability of solutes and water in epithelia and endothelia.^12–14^ In addition, non-junctional claudins can serve as CPE receptors.^15–17^ Within a short time after receptor-binding, severe damage to epithelial cells occurs as due to formation of cytotoxic pores in the plasma membrane^18–21^. Ca^2+^ influx triggers cell death by oncosis or apoptosis. ^22–24^

CPE functions as pore-forming toxin (PFT), which relies on oligomerization on cell membranes and insertion of a membrane-spanning segment. CPE was suggested to belong to the subgroup of β-PFTs that form a β-barrel pore.^25^ β-PFTs include MACPF/CDC, ABC toxins, aegerolysins, colicins, AB toxins, and the emerging superfamily of aerolysin/ETX-MTX-2 proteins.^26^ The N-terminal domain of monomeric CPE (1-205) shows structural homology with the C-terminal pore-forming domain of aerolysin, thus CPE was assigned to the aerolysin/ETX-MTX-2 superfamily.^27^

The soluble CPE monomer (35 kDa) is organized in two main functional domains: (1) the N-terminal domain (nCPE, residues 1-202), which facilitates the toxin’s cytotoxicity by pore formation, and (2) the C-terminal domain (cCPE, residues 203–319), which binds to the claudin receptor on target cells. The crystal structures revealed that cCPE is made up from one (I) and nCPE from two structural domains (II and III). Each of these domains consists of one α-helix and several, mostly antiparallel β-strands (I: 9 C; II: 5 β; III: 3 β; **Figure S1A**).^27, 28^

Biochemical and structural data revealed that in particular the residues Y306, Y310, Y312 and L315, located at the base of a binding pocket of cCPE, are responsible for the nanomolar affinity to certain claudins, namely CLDN3, -4, -6, -7, -8, -9, -14, and -19.^29–31^ On the receptor site, the claudin’s ECS, which form an antiparallel β-sheet composed of five β-strands, are involved in the interaction. The ECS2 portion, consisting of strand β5 and an elongation of TM3, has been shown to bind in the described pocket in cCPE.^32–35^ The strong interaction with CPE is mediated by the CLDN-ECS2 motif (N/D(P-1) P/S(P) L/M/V/I/S(P+1) V/T(P+2) P/A/N/D(P+3)), letters indicate amino acids, and P refers to conserved proline in ECS2, e.g. P150 in CLDN4 (**Figure S1D**).The more stringent motif (D/E(P-4) x(P-3) x(P-2) N(P-1)P(P) L/M/V(P+1) V(P+2) P/A(P+3) was identified for high-affinity interaction.^15, 36–38^ Moreover, the β1-β2 loop of ECS1 contributes to the binding.^33–35^

Upon binding, cCPE structure remains unchanged, while according to biochemical data, a non-cytotoxic "small complex" with a molecular weight of ∼90 kDa is formed, consisting of CPE, a receptor claudin and possibly a non-receptor claudin.^39^ It was further suggested that several small complexes oligomerize on the cell surface before pore formation.^40^. Homology of the CPE monomer to a HA3 botulinum toxin component and continuous studies of stoichiometry and mass of the developing bigger prepore or pore complex lead to the hypothesis that six small complexes are involved.^27, 39^ Therefore, the resulting 425–500 kDa complex is also referred to as "CPE hexamer-1" (CH-1). In mammals, a further heavier, 550–660 kDa CH-2 complex was found, which includes occludin, another TJ protein.^39^ In contrast to cCPE, the two structural domains of nCPE are proposed to undergo drastic conformational changes for the subsequent cytotoxic pore formation.^27^ A stretch of amphipathic residues (residues 81-106) in Domain II has been suggested as the membrane-spanning region.^40^ However, this could not yet be proofed by means of a 3D structure of the pore complex. Due to the drastic conformational change in the N-terminal domain, existing crystal structures of the CPE monomer^27, 28^, CLDN3^41^ and CLDN4^34^, provide insufficient information about the formation and structure of the final pore. Here, in line with the existing biochemical data, based on AlphaFold2 and molecular dynamics (MD) simulations, we propose a molecular model for the prepore as well as the formation of the final pore complex.

## Results

### Alphafold2-based prediction of CPE oligomerization

Several structures of cCPE bound to CLDN3, -4, -9 or -19 (PDB IDs 3X29, 5B2G, 6AKE, 6OV3, 7KP4, 8U4V); ^33–35, 41–43^, and of claudin-free CPE (PDB IDs 2XH6, 3AM2, 3ZIX, 8U5F; ^27, 28, 44, 45^ have been resolved (**Figure S1**). These, along with comparisons to other β-PFTs with known pore structures (PDB IDs e.g. 3J9C, 5JZT, 5GAQ; ^46–48^, strongly indicate that CPE undergoes major conformational changes during pore formation. However, these changes remain unclear because no CPE pore structures, either alone or in complex with claudins, currently exist.

Given the advances in prediction of protein complexes by AlphaFold2, we used its variant ColabFold v1.5.5^49, 50^ for prediction of CPE homo-oligomers. Several input parameters, including full-length CPE (1-319) and truncated sequences, were used (see Methods for details). Only prediction results consistent with experimental key findings were considered, specifically hexa- or heptameric ring-shaped architectures with a central pore formed by nCPE. Initially, the most meaningful results were obtained with CPE_1-202_ (nCPE), which lacks the claudin-binding domain (cCPE, 203-319). We conclude that the removal of cCPE in the prediction suppressed the over-contribution of non-pore-forming folds from full-length monomeric CPE structures and cCPE/claudin complex structures (PDB IDs listed above) in the AlphaFold training data sets. Some of the output CPE oligomer structures (e.g., state nCPE-IV in **Figure 1E-G**) reproduced the fold of nCPE found in the CPE crystal structures (PDBs 2XH6, 3AM2, 3ZIX, 8U5F; **Figure S1**), while others showed a different conformation for the region 70-116. Importantly, the latter oligomeric structures (states nCPE-V to –VII, **Figure 1H-M**) and the aforementioned one (state nCPE-IV, with nCPE fold of crystal structures, **Figure 1**) contained a central pore-shaped β-barrel. Strikingly, the length of the barrels increased from state nCPE-IV to nCPE-VII (30 Å, 34 Å, 53 Å and 86 Å, respectively), suggesting subsequent conformational states converting the initial, receptor-unbound monomeric CPE fold via pre-pore intermediates into the functional membrane-permeating pore. Thus, we proceeded with a step-by-step analysis of the ColabFold models nCPE-IV to VII.

**Figure 1:**
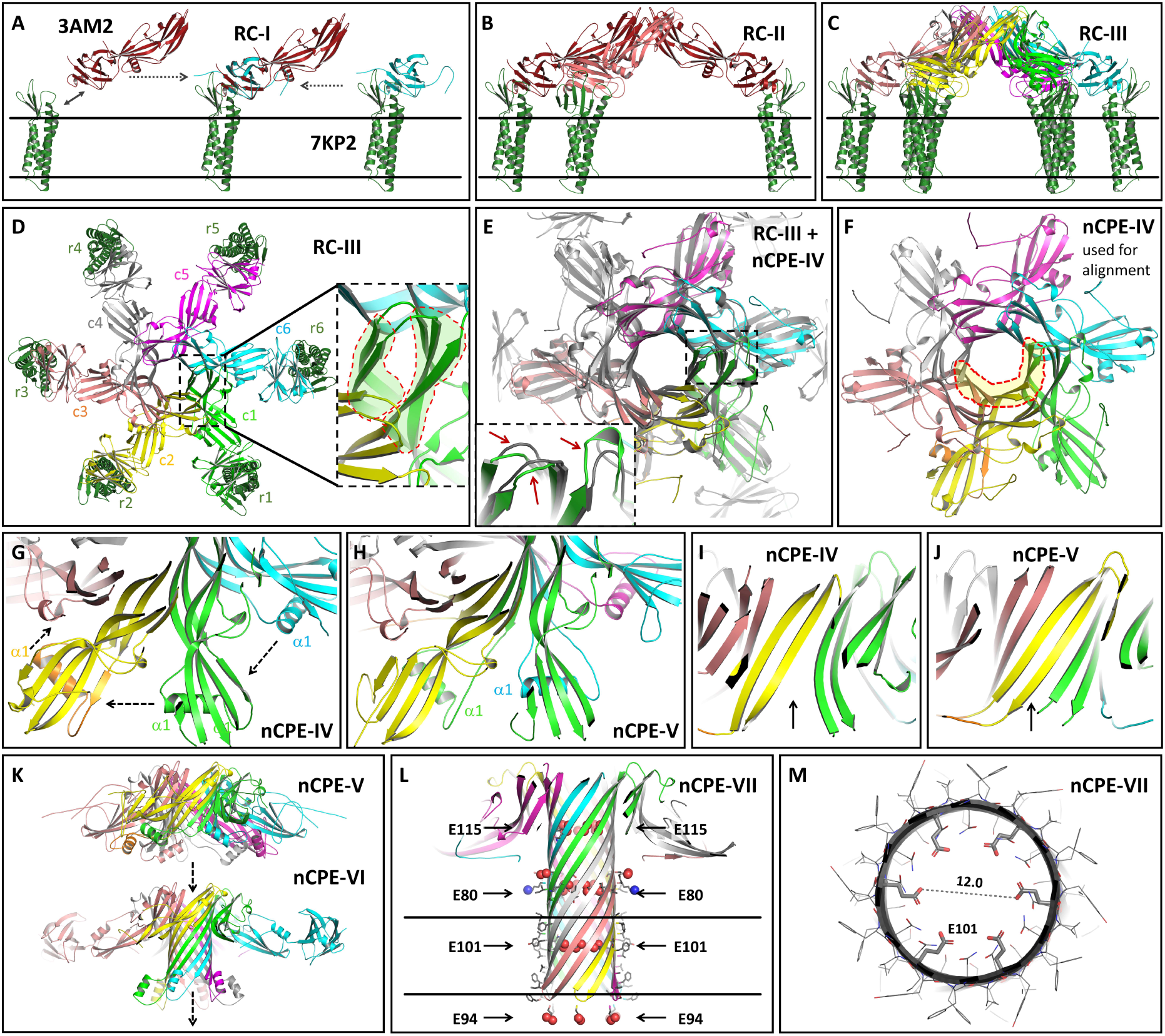
Use of Colabfold/Alphafold2 and structural alignment with CPE and cCPE-CLDN4 complex crystal structures (PDB 3AM2 and 7KP4) enables modeling of CPE pore complex and assembly intermediates. **(A)** Binding of monomeric CPE to CLDN4 in membrane leads to state RC-I. **(B-D)** Oligomerization into state RC-III. (**C**) Side view of RC-III. (**D**) Top view of RC-III. CPE (c1-c6) and claudin receptor (r1-r6) subunits are labeled. Insert: β3-& β5-strands forming inner and β2-, -β6-& β7-strands forming outer β-barrel are shown (marked area). (**E**) Superposition of states RC-III (gray) and nCPE-IV (colored). Insert: To avoid minor clashes, nCPE-IV (green) differs slightly from RC-III (gray) as highlighted by none overlapping protein cartoons (red arrows). **(F)** State nCPE-IV used for alignment of the other states. Inner β-barrel fit is marked. (**G-J**) nCPE-IV to nCPE-V transition. (**G, H**) Direction of movement/swapping of α1-region between CPE subunits indicated by arrows. (**I, J**) Improvement of β-barrel indicated by arrows. **(K, L)** Stepwise β-barrel extension in states nCPE-V, nCPE-VI and nCPE-VII. (**L**) Three hexa-glutamate rings (E80, E101 and E115) forming constrictions along the β-barrel pore are indicated by arrows. Also, hexa-glutamate ring formed by E94 at bottom opening is shown. Negatively (red) and positively (blue) charged groups shown as spheres, hydrophobic residues facing lipids shown as sticks, membrane boundaries indicated (black lines). (**M**) Ring formed by E101 in middle of transmembrane region. Clipped view, E101 as sticks, other residues as lines, distance between E101 O atoms.

### Decoding CPE Pore Formation: Insights from alignment of Alphafold2-predicted oligomers with solved CPE and cCPE/CLDN4 structures

To connect the selected Alphafold2 predictions with the experimentally solved CPE/claudin structures, we first aligned all structures to the one with the highest segment overlap with all individual structures, which is the conformational state nCPE-IV (**Figure 1F**). This alignment allowed us to develop a sequential model of pore formation (**Figure 1A-L**). Initially, monomeric CPE binds to a receptor such as CLDN4 in the cell membrane (**Figure 1A**). This conformation, referred to as state receptor-CPE I (RC-I) of the CPE/claudin complex, was generated by aligning the structure of CPE (PDB ID 3AM2) with a complex structure of cCPE bound to human CLDN4 (PDB ID 7KP4), combined with a membrane embedment prediction for CLDN4 (see methods). According to gel shift analysis and other biochemical data ^39^, six CPE/CLDN4 complexes oligomerize via nCPE (**Figure 1B-D**, states RC-II and RC-III). State RC-III exhibits cyclic symmetry and contains a short β-barrel-like pore formed by two β-strands (residues 68-77 (β3) and 111-120 (β5) from each of the six CPE subunits in the center, surrounded by a second, looser β-barrel formed by three additional β-strands of the CPE subunits (residues 57-63 (β2), 127-135 (β6), 158-167 (β7) (**Figure 1D**, inset). This architecture resembles the central cap of the mushroom-like cap-and-stem architecture seen in other β-PFTs (PDB IDs e.g. 3J9C, 5JZT, 5GAQ; ^46–48^). Importantly, the nCPE segments of state RC-III align well with the nCPE subunits of the Alphafold2-predicted state nCPE-IV (Cα RMSD of ∼1.9 Å, **Figure 1E, F**). The main difference between nCPE in state RC-III (manual assembly) and nCPE-IV (AlphaFold2 prediction) is that minor steric clashes in RC-III (**Figure 1E**, inset), which result from the structural alignment of six rigid CPE subunits to the state nCPE-IV structure, are absent in nCPE-IV. This state likely represents a CPE pre-pore structure formed by initial CPE oligomerization and not yet penetrating the membrane, similar as shown for state RC-III in **Figure 1C**.

Interestingly, another AlphaFold2-predicted structure (state nCPE-V) is very similar to state nCPE-IV, with the difference that it contains a notable domain swap of the amphipathic α-helix segment α1 (residues 92-105) in domain II (**Figure S1A**). In state nCPE-IV, it interacts with the β-sheet (β1, 4, 6, 7, 8) in the CPE domain II (**Figure S1A**) of the same subunit. However, in state nCPE-V, it interacts with the corresponding β-sheet of a neighboring CPE subunit (**Figure 1G, H**, **Movie S1**). This swap closes a gap in the β-barrel lining of the pre-pore (**Figure 1I, J**). In the next modeled state nCPE-VI, the interaction of the α1-segment (92-105) with the β-sheet in CPE domain II is lost and the adjacent segments 77-83 and 106-111 extend the central β-barrel, resulting in its elongation from 33 to 53 Å (**Figure 1K**). Finally, the region 84-105 stretches into a β-hairpin and extends the β-barrel to the full stem (86 Å) of the mushroom-like architecture and forms the membrane-spanning region (**Figure 1L**, state nCPE-VII, **Movie S2, S3**). Importantly, in the region 81-108, hydrophobic residues are pointing outwards whereas hydrophilic residues are pointing towards the pore lumen. Beyond the region 81-108, hydrophilic residues face outwards. Along the β-barrel pore lumen, three hexa-glutamate rings are formed by E101, E80, E115, respectively. The O-atoms of the glutamate residues in the three rings are ∼12 Å apart from each other (**Figure 1L, M**). Therefore, the pore lining fits perfectly to a cation-selective transmembrane pore formed by CPE, which brings the model in agreement with conductivity and other experiments.^51, 52^

### Comparison of hexamer and heptamer pore models of nCPE with structures of other β-pore forming toxins

Monomeric CPE shows structural homology to *A. hydrophila* aerolysin that forms a heptameric pore. ^27, 28^ Hence, we also predicted nCPE heptamers with Alphafold2. Some of these predictions resulted in a heptameric pore with a similar stem and cap center architecture as that of the aerolysin pore (**Figure S2**). However, when compared to the nCPE hexamer model, the heptameric pore had the following inconsistencies: (i) The β-hairpin tip was irregular and the length of the pore barrel was shorter for the heptamer (78 Å instead of 86 Å), impairing it from fully spanning the membrane (**Figure S2A, B**; see also below); (ii) the hydrophilic Q97 that points towards the pore lumen in the hexamer points towards the hydrophobic lipids surrounding the pore in the heptamer. Conversely, the opposite was true for the hydrophobic I96 (**Figure S2D, E**). For the heptamer, these residue orientations would be energetically unfavorable. Due to these observations and the biochemical data supporting a CPE hexamer^39^, the heptamer model was not analyzed further.

Comparison of the nCPE hexamer pore model (state nCPE-VII) with the structures of the other β-PFTs *A. hydrophila* aerolysin, *C. perfringens* epsilon toxin (ETX) and *E. fetida* lysenin showed that all these toxins share a similar mushroom-like architecture with a β-barrel pore stem and two concentric β-barrels in the center of the cap region (**Figure 2**). However, while aerolysin and ETX form heptameric pores and lysenin forms nonameric pores, CPE forms a hexamer. The nCPE model shares this hexameric architecture, and also structure and sequence homology with the HA3 subcomponent of the hetero-hexameric botulinum progenitor toxin (sequence identity: 27% between CPE1-200 and HA3a_8-184_ and 25% between CPE_1-319_ and two of three domains in HA3b.^27^ The corresponding crystal structure (**Figure 2D-F**; PDB ID 2ZS6,^53^) was one of the templates used by AlphaFold2 for the predictions (see methods). Similar to the other toxins, HA3 contains two concentric β-barrels surrounding the central pore. In contrast to the CPE model, the HA3 pore is of insufficient length to cross the membrane, thereby likely resembling an inactive prepore state (**Figure 2D-F**). Consistently, the HA3 oligomer structure shows most structural similarity with state nCPE-IV with which it also shares the 5-stranded β sheet (β1, 4, 6, 7, 8) and the interacting α-helix (α1 of the CPE domain II (**Figure 1F, 2E, S1A**).

**Figure 2:**
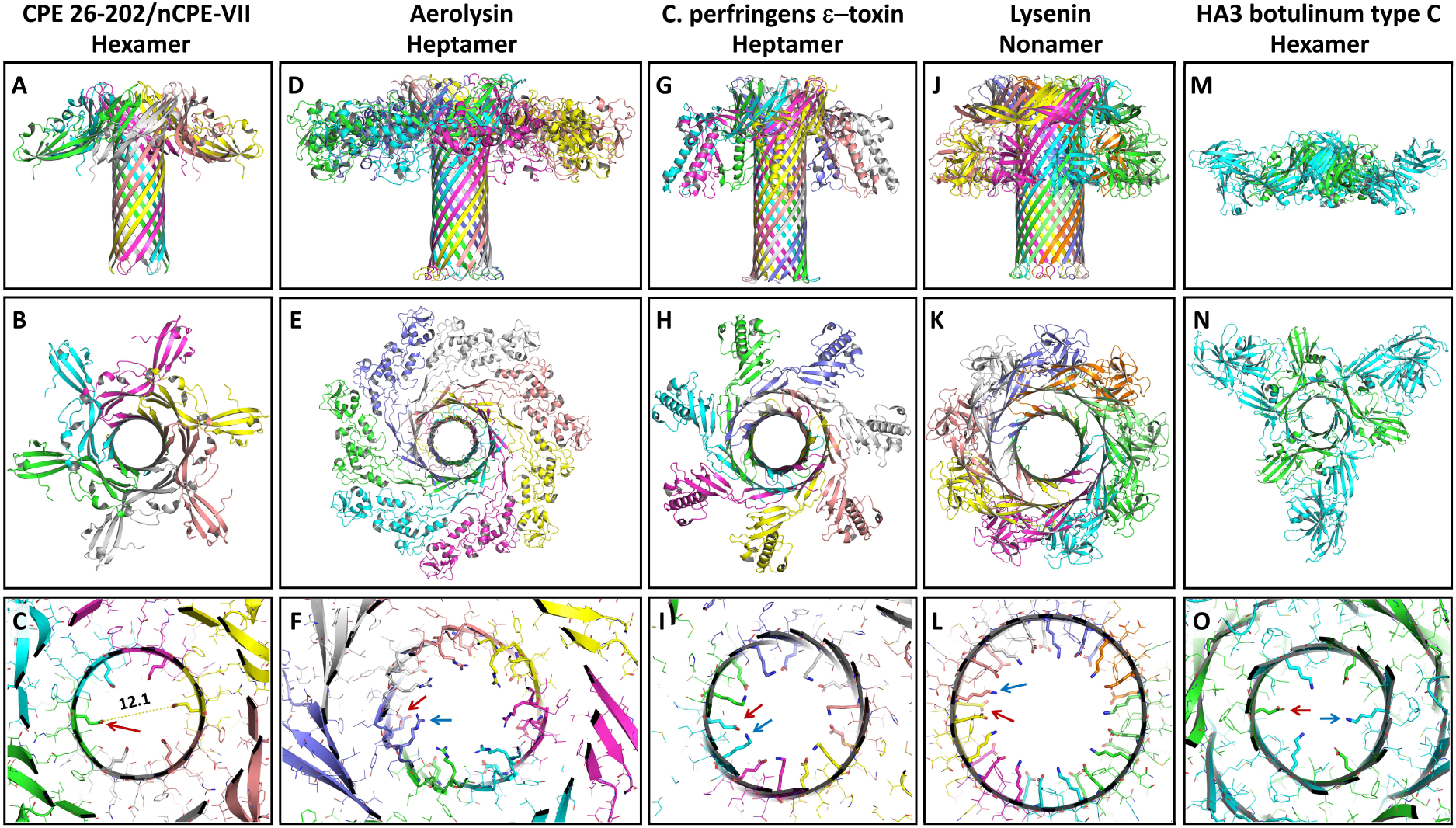
Comparison of the nCPE hexamer model (**A-C**) with the β-pore forming toxins aerolysin, PDB 5JZT, homo-heptamer (**D-F**), *C. perfringens* ε-toxin, PDB 6RB9, homo-heptamer (**G-I**), lysenin, PDB 5GAQ, homo-nonamer (**J-L**) and HA3 botulinum type C, PDB 2ZS6, hetero-hexamer (**M-O**). Upper panels, side view; middle panels, top view; lower panels, close up of pores in cap region of the toxin complexes with charged residues shown as sticks (arrows) and other residues as lines, O-atoms in red and N-atoms in blue.

The diameters of the different toxins increase with the number of subunits. In addition, the pore lining differs strongly. While CPE has numerous negatively charged residues but lacks positively charged ones in the pore lumen, the pores of other toxins contain both (**Figure 2**, lower panels). This charge distribution fits to the CPE-specific cation selectivity, which is not observed for the other β-PFTs.^52^ In summary, the observed similarities and differences between the presented experimental structures and the hexameric nCPE pore model support the plausibility of the latter.

### Conformational change of the cCPE-nCPE linker region during transmembrane pore formation

Next, we examined the anchoring of the hexameric CPE pore to the claudin receptors. For this, state nCPE-VII was aligned with state RC-III (via a common alignment to state nCPE-IV, **Figure 1C-L**). The membrane plane was predicted for the 6 claudin subunits using the PPM server ^54^; https://opm.phar.umich.edu/ppm_server3_cgopm). In the resulting complex (state RC-VII) consisting of 6 CLDN4, 6 cCPE and 6 nCPE chains, the tips of the β-hairpins of the pore did not entirely span through the membrane (∼13 Å missing, **Figure 3A**, state RC-VII). As a potential mechanism that enables a transmembrane pore, we speculated a conformational change in the linker region between the cCPE and the nCPE domains (residues K193-A205, blue loop segment in **Figure 3, D-G**). Our predicted pore structure (RC-VII, **Figure 3C**) did not yet contain a linker between nCPE (from nCPE-VII) and cCPE (from 7KP4 of RC-III). For linkage and to move nCPE but not cCPE downwards (relative to CPE in RC-III) as a rigid body, most of the linker region (K197-A205, part of blue region in Figure 3D-G)) was manually joined to cCPE. For this, H146 at the tip of nCPE was moved from L278 at the upper rim of cCPE to the lower rim close to W234 (**Figure 3B, D-G**, black dashed arrows). This resulted in a full-length CPE hexamer pore (state RC-VIIIa), in which, unlike as in state RC-VII, the β-hairpin tips of the pore reach the lipid carbonyls on the cytoplasmic side of the membrane (**Figure 3A, B, E**).

**Figure 3:**
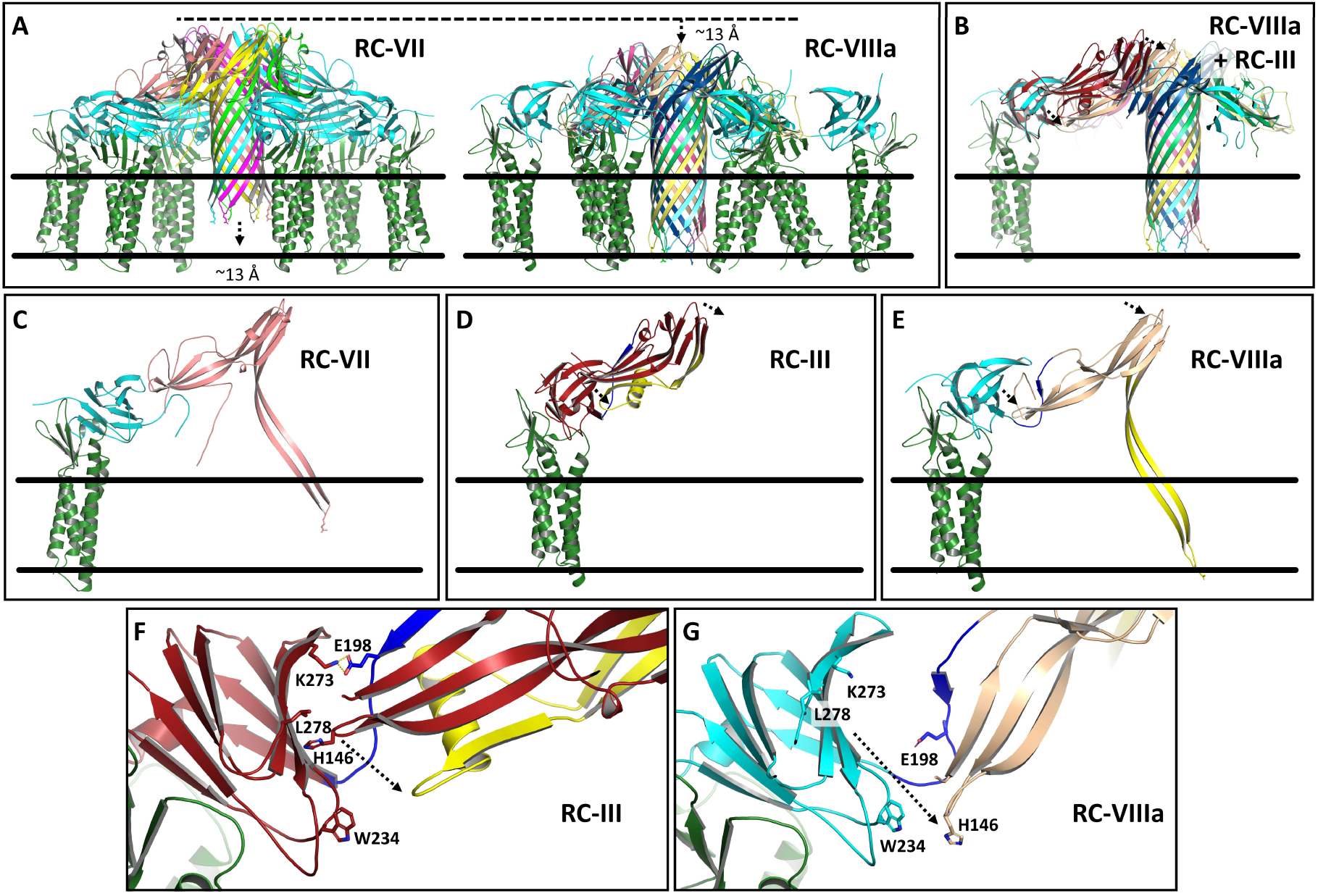
Conformation change in linker region between cCPE and nCPE (residues 193-205, blue) allowing full transmembrane penetration of β-hairpin tips of pore. (**A**) Comparison of CLDN4-bound CPE hexamer pore complexes without (state RC-VII) and with downwards shift of nCPE (state RC-VIIIa). E94 at β-hairpin tip is shown as stick. (**B**) Clipped view of state RC-VIIIa focused on one claudin with superimposed state RC-III to visualized the conformational difference between the monomeric CPE structure (3AM2 of RC-III, red) and CPE in RC-VIIIA (cyan, beige). Individual CPE/claudin dimers are shown separately for RC-VII (**C**), RC-III (**D**) and RC-VIIIa (**E**). The region 73-116 that changes conformation is labeled in yellow. In RC-III it includes the α1-helix, in RC-VII and RC-VIIIa it forms the main part of the pore β-barrel. The manual shift of nCPE in RC-VIIIa relative to the position in RC-III is labeled by dashed arrows in (B, D-G). (**F**, **G**) Close-ups of (D, E) highlighting the move of H146 at tip of nCPE from close to L278 of cCPE in RC-III towards W234 of cCPE in RC-VIIIa.

We also tested another cCPE-nCPE linker variant (state RC-VIIIb) based on an AlphaFold2 prediction for CPE26-319 (see methods). Similar as for state RC-VIIIa, nCPE was shifted downwards so that H146 of nCPE was close to P233 at the lower rim of cCPE (**Figure S3H-J**, black dashed arrow). This resulted in pore state RC-VIIIb in which the β-hairpins of the pore were also moved towards the cytoplasmic side of the membrane (**Figure S3**). However, the conformation in the region P191-A205 (blue in **Figure 3D-G, S3H-J**) was with a rather unlikely β-to-α transition for RC-VIIIb stronger changed than for RC-VIIIa with respect to the CPE subunit (PBD 3AM2, **Figure S3**). Thus, we concentrated mainly on state RC-VIIIa for further analysis.

### MD simulations confirm the modeled mushroom-like architecture and claudin anchoring of the CPE pore

The pore model states nCPE-VII, and RC-VIIIa were further analyzed with respect to their stability and ion permeability using MD simulations. For this, a Schrödinger Maestro- and Desmond-based platform and workflow was used as previously established for simulating claudin-based paracellular ion channels.^55, 56^ The above-mentioned protein complexes were embedded in POPC lipids. NaCl and water were added, the system was relaxed, stepwise equilibrated, and finally simulated without constrains for 100 ns using the OPLS4 force field.

First, the nCPE hexamer state nCPE-VII was subjected to simulations. The whole stem and cap structure remained stable throughout the simulation, and the proposed transmembrane part of the stem (residues 80-108) remained embedded in the membrane (**Movie S4**). This included the β-hairpins, whose negatively charged tip (E94) reached to the cytoplasmic side, all along the water and ion filled pore barrel and the two concentric β-barrel elements in the cap region (**Figure 4A**). Preservation of the structural architecture throughout simulation is reflected by a RMSD of ∼3.85 Å (protein backbone, mean RMSD for last 50 ns of simulation with respect to initial structure). The three hexa-glutamate rings formed along the pore by E101, E80, and E115, respectively, interacted strongly with Na^+^ ions (**Figure 4B-E**). Only very few Cl^-^ ions were observed within the pore. Thus, the simulation result was plausible.

**Figure 4:**
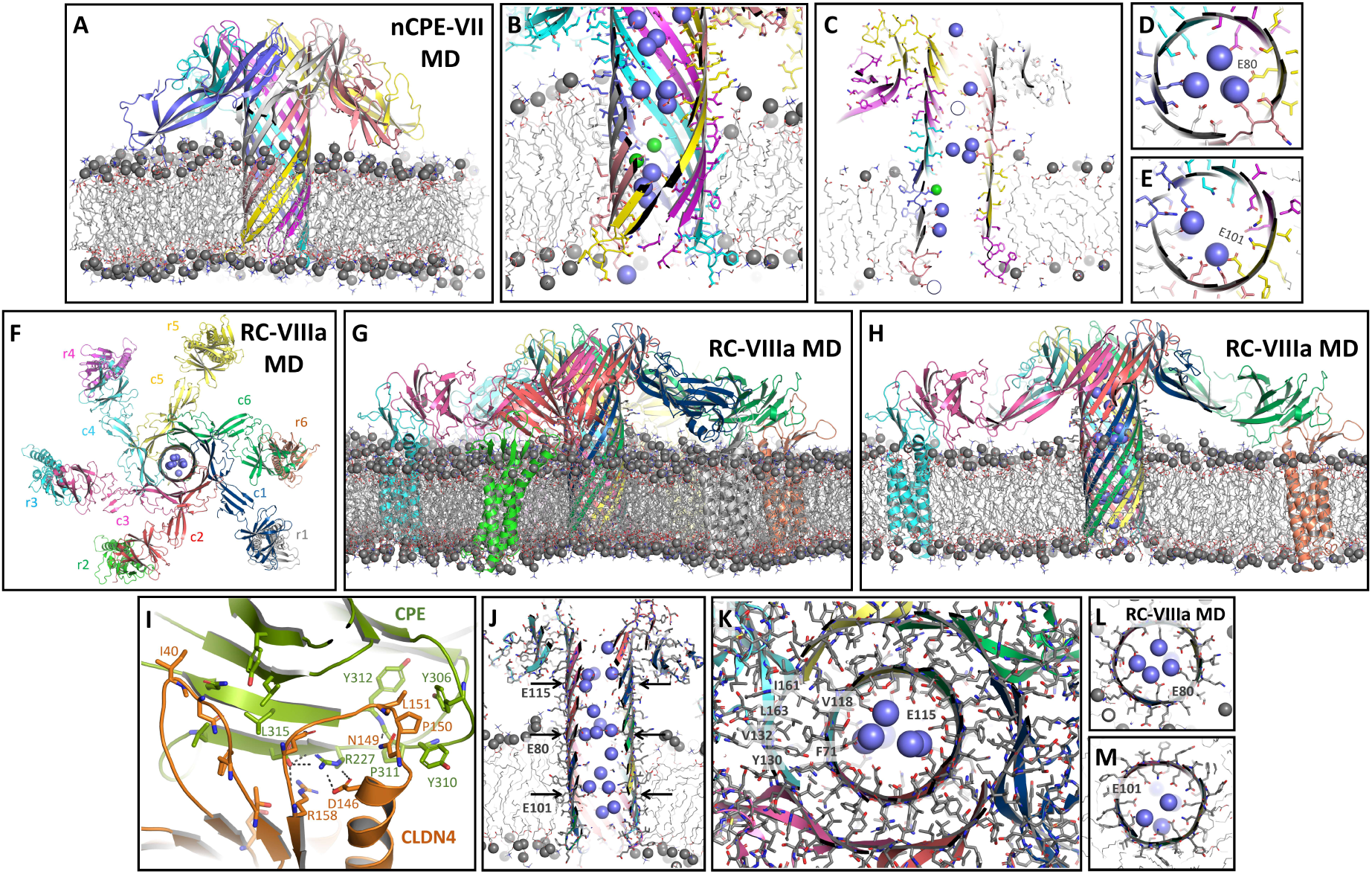
MD simulations of hexamers of nCPE (nCPE-VII, **A-E**) and Cldn4/CPE complexes (RC-VIIIa, **F-M**). Shown are snapshots after 100 ns of free simulation with protein as cartoon, relevant residues as sticks, lipid acyl chains as gray lines, phosphate head groups as gray spheres, sodium ions as blue and chloride ions as green spheres. (**A**) For nCPE-VII the pore barrel outside and inside the membrane, and the pore cap are well preserved (∼3.85 Å mean RMSD for last 50 ns of simulation with respect to initial structure, protein backbone). The pore is well embedded in the membrane. (**B, C**) The pore is clipped in two versions to illustrate membrane embedment and strong presence of sodium ions in the pore. Rings formed by six E80 residues slightly above membrane plane (**D**) and by six E101 residues within the membrane plane (**E**) strongly attract cations. (**F**) RC-VIIIa shown as top view with labeled CPE (c1-c6) and CLDN4 receptor (r1-r6) subunits. (**G, H**) RC-VIIIa shown as side view (clipped in (H)). The pore barrel and the pore cap are well preserved (4.57±0.37 Å mean RMSD for last 50 ns of simulation with respect to initial structure, protein backbone). The pore complex is well embedded in the membrane. (**I**) Details of the stable interaction between cCPE domain and CLDN4. Key residues are labeled and key electrostatic interaction shown as dashed lines. (**J**) Clipped side view of pore to illustrate membrane embedment and strong presence of sodium ions in the pore lumen. (**K**) The inner and outer β-barrel in the cap region (upper arrow in (J)) are held together mainly by hydrophobic interactions. Some key residues are labeled. In the center, ring of six E115 residues strongly attracts cations (spheres). Rings formed by six E80 residues (**L**) slightly above membrane plane (middle arrow in (J)) and by six E101 residues (lower arrow in (J)) within the membrane plane (**M**) also strongly attract cations.

Next, MD simulations were performed for systems containing the full-length CPE hexamer bound to six CLDN4 receptor subunits (state RC-VIIIa, Figure 4F-M). The transmembrane regions of both the CPE hexamer (residues 80-108) and the six claudins were well embedded in the membrane, and the whole complex remained stable throughout the simulation (4.57±0.37 Å, mean RMSD for last 50 ns of simulation with respect to initial structure, **Movie S5**). This included the β-hairpins, their cytoplasmic tips (E94), the inside and outside of the stem barrel, and the concentric β-barrel element in the cap region (**Figure 4F, G, H, J**), similar as for the simulation of state nCPE-VII (**Figure 4A-C**). In addition, the interaction between cCPE and CLDN4 remained stable throughout the simulation, including the interfaces observed in the experimental cCPE/CLDN4 structures (5B2G, 7KP4, 8U4V,^34, 43, 57^). In particular, L151 in the ECS2 of CLDN4 was stably plugging into the triple tyrosine pocket (306, 310, 312) of cCPE (Figure 4I, ^32, 58^). Details are shown below for a subsequent simulation (F**igure 8**, **9**). The cap region was stabilized by the two concentric β-barrels (**Figure 4K**). Here, the intramolecular contact between two β-sheets is maintained mainly by hydrophobic interactions (between V58, L63, F71 and V118 of the inner barrel and T120, V128, Y130, V132, I161, L163 of the outer barrel), similar as in the monomeric CPE (PBD 3AM2). Inter-protomer interactions are formed by H-bonds between neighboring β-strand and by electrostatic side chain interactions (e.g. E67-K165). The three hexa-glutamate rings formed along the pore by E101, E80, E115, respectively, interacted strongly with Na^+^ ions (**Figure 4J-L**). Similar results were obtained for the respective simulation of CPE hexamer state VIIIb (**Figure S4**). In summary, the MD simulations strongly supported the modeled architecture, claudin anchoring, membrane embedment, pore lining, and cation selectivity of the CPE hexamer pore.

### Do claudins form a dodecameric ring during CPE pore formation?

Biochemical data suggest that the CPE pore complex contains more claudin than CPE subunits.^39^ Furthermore, in CPE complexes, low affinity CPE receptors such as CLDN1 or CLDN5 have been detected in addition to high affinity CPE-receptors such as CLDN4 and CLDN3 ^39, 59^). Molecular modeling^32^ predicted and crystal structures^34^ verified a 1:1 stoichiometry of the high affinity cCPE-claudin interaction. In contrast, for the CPE-claudin complex, a 1:2 stoichiometry was suggested. ^39, 60^ Thus, either only one subunit of a claudin dimer interacts directly with CPE, or the second one is in contact with cCPE and/or nCPE domains via additional low-affinity interaction sites. For the second claudin within the complex, different binding configurations are conceivable. However, due to the polymerization capability of claudins, we speculated that 12 claudin subunits could form a dodecameric ring via a bended variation of the linear cis-interaction found in polymeric claudin-based tight junction strands. ^14, 55, 61–63^ It was shown that binding of cCPE to claudins alters the conformation of the claudin regions involved in the linear cis-interaction.^41, 64^ In addition, it was suggested that differences in these interface regions can affect the rotational angle or relative positions between/of neighboring claudin subunits.^41, 56, 63^ Thus, it is conceivable that binding of cCPE to claudins bends the otherwise rather straight cis-interface by up to ∼30° leading in sum to a 360° rotation for a dodecamer and consequently to the formation of a claudin ring. This putative ring was modeled on the base of a linear-cis interface in claudin strand (**Figure S5H**) and 30° rotations between the subunits (**Figure 5, Figure S5** and see methods). The ring diameter was smaller than that of the discontinuous ring formed by the six claudins in the complex state RC-VIIIa (∼148 Å vs ∼182 Å for S58-S58 Cα distance, **Figure S5A** and **Figure 5D**). Consequently, for docking CPE to the ring, the straight connection between nCPE and cCPE of state RC-VIIIa had to be modified to an angled one, in order to keep the structural integrity of the nCPE hexamer and that of the cCPE-CLDN4 complex. This resulted in a tightly packed and sterically fitting complex (state RC-IXa) without steric conflicts in which each second CLDN4 subunit interacted with CPE as in state RC-VIIIa. The other claudin ring subunits were in loose contact on one side with cCPE and on the other side with the domain II of nCPE (**Figure 5**). The full conformational change of CPE from the monomeric state RC-I to the final state RC-IXa is depicted in **Figure 6**. In addition, we generated a variation of state RC-IXa (state RC-IXb) in which the tilt of the claudins in the membrane plane was slightly different. Details of this variant are shown in the supplement (**Figure S6**).

**Figure 5:**
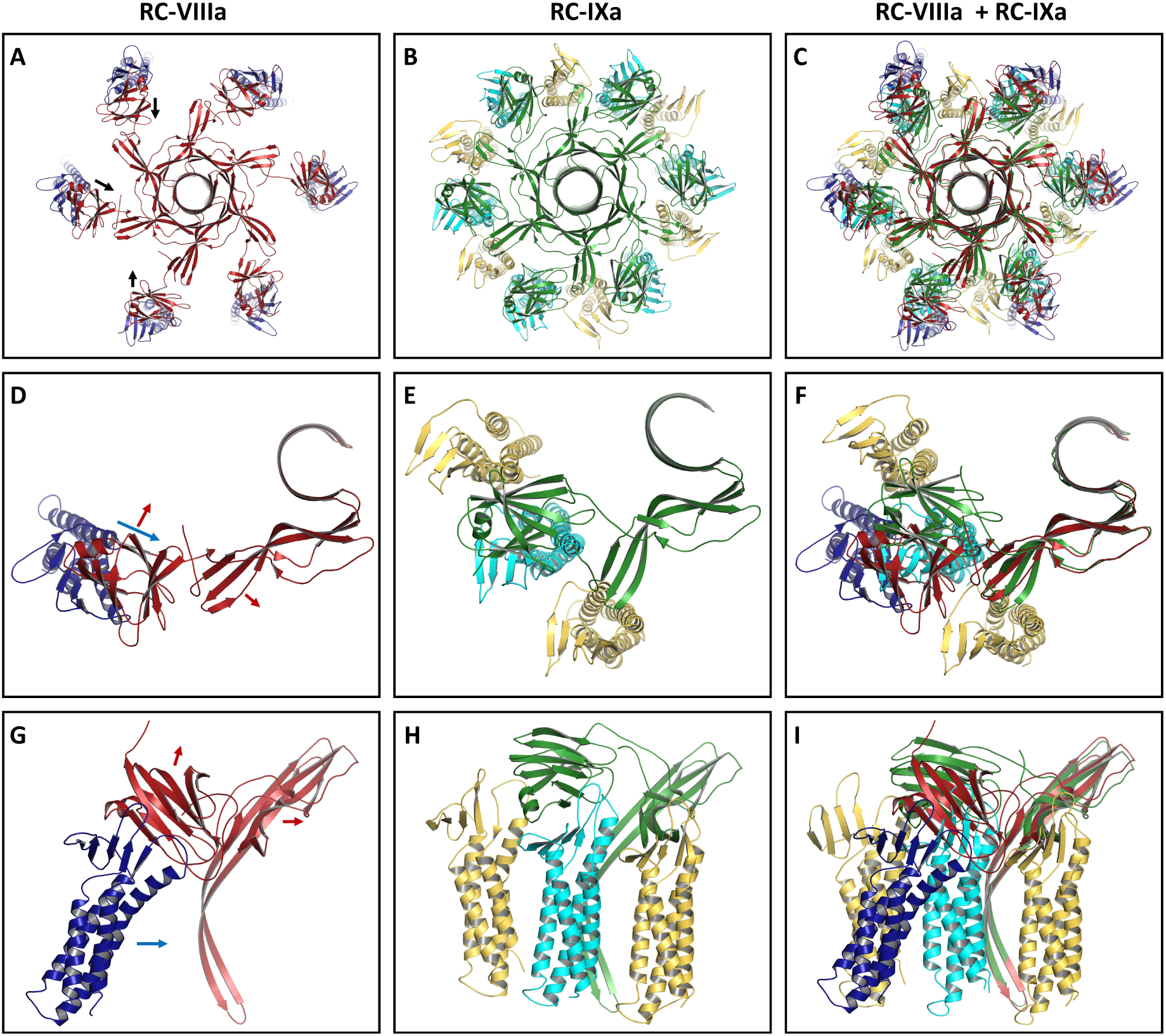
Details of modification of relative cCPE/nCPE positions for state RC-IXa. (**A-C**) Top views. (**A**) State RC-VIIIa: CPE pore hexamer (red) bound to six individual claudins (blue). (**B**) State RC-IXa: CPE pore hexamer (green) bound to a dodecameric claudin ring. Claudin subunits primarily bound to CPE are colored cyan, additional claudin subunits are colored yellow. (**C**) Superposition of RC-VIIIa and RC-IXa. (**D-I**) Single CPE subunits bound to claudins are shown in top view (**D**, RC-VIIIa), (**E**, RC-IXa), (**F**, both) or side view (**G**, RC-VIIIa), (**H**, RC-IXa), (**I**, both). For RC-VIIIa, one CPE subunit bound one claudin subunit. For RC-IXa, in addition to the primarily CPE-bound claudin subunit, both adjacent claudins are shown. The arrows indicate direction of movement of cCPE (red) and claudin (blue) from state RC-VIIIa to state RC-IXa. 100 ns snapshots of MD simulation are shown.

**Figure 6:**
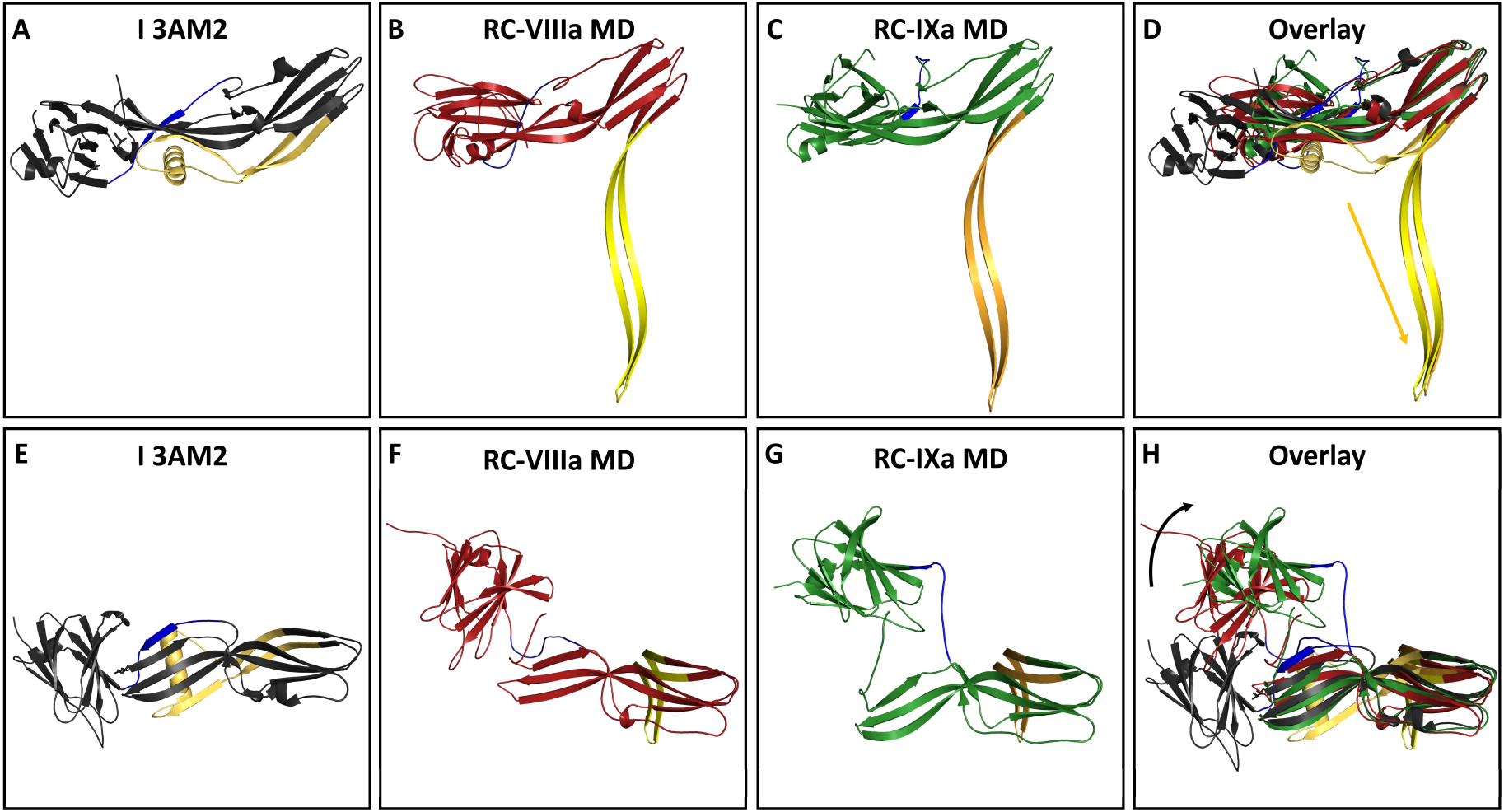
Comparison of CPE monomer crystal structure (**A**, side view, **E**, top view) with state RC-VIIIa (**B**, side view, **F**, top view) and state RC-IXa (**C**, side view, **G**, top view) and superposition of the three structures (**D**, side view, **H**, top view). Conformational changes: Yellow: residues 70 -116, region forming the pore barrel (straight arrow); blue: 191-204, linker between cCPE and nCPE. Note the shift and turn of cCPE relative to nCPE (round arrow) between the states. For RC-VIIIa and RC-IXa snapshots after 100 ns MD simulation are shown.

### MD simulations support a cation-selective CPE pore anchored to a dodecameric claudin ring

The pore complex states RC-IXa and RC-IXb, each containing six CPE and twelve CLDN4 subunits, were embedded in a membrane, and the two respective MD simulations were performed as done for other complex states. Both complexes showed a similar stability with respect to the pore stem, the cap region, the main cCPE-claudin interaction site, membrane spanning, pore lining, and cation attraction as the simulations of the complex states nCPE-VII to RC-VIIIa (**Figure 7, S7, S8, Movie S6**). The following final and detailed analysis of MD simulations performed in this study was focused on the simulation of state RC-IXa, since the ones with the dodecameric CLDN4 ring appeared to be the most informative and complete states. Also, compared to state RC-IXb, the tilt of the claudins within the membrane in state RC-IXa agreed with the tilt observed in tight junction strand and channel models.^55, 56, 62^

**Figure 7:**
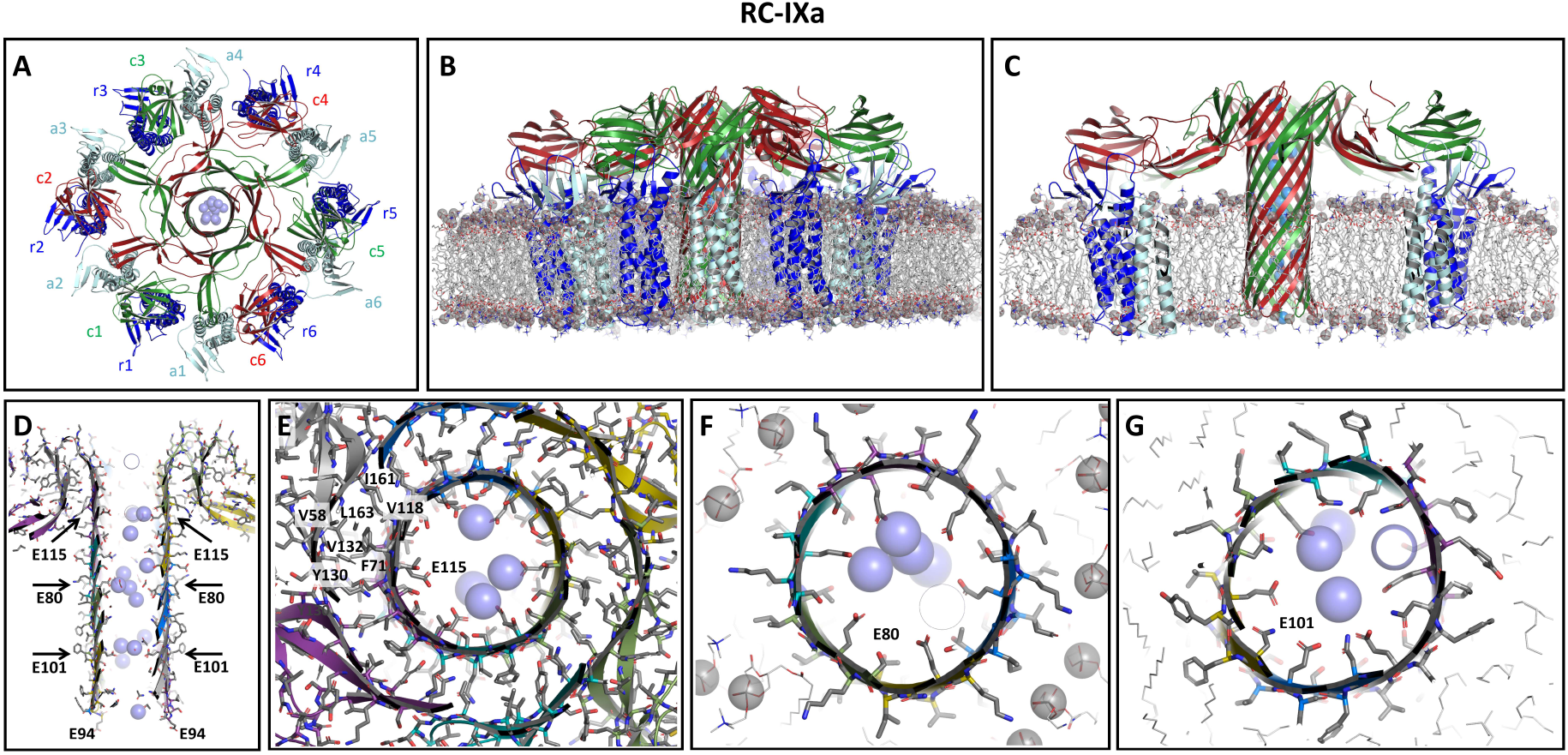
MD simulation of CPE hexamer anchored to dodecameric CLDN4 ring (pore complex state RC-IXa). Shown are snapshots after 100 ns of free simulation with protein as cartoon, relevant residues as sticks, lipids acyl chains as gray lines and phosphate head groups as gray spheres, sodium ions as blue spheres. **(A)** Top view with CPE (c1-c6, green & red), canonical receptor CLDN4 (r1-r6, blue) and additional CLDN4 (a1-a6, cyan) subunits. (**B, C**) Side view (clipped in (C)). The pore barrel and the pore cap are well preserved. The pore complex is well embedded in the membrane. (**D**) Clipped side view of pore to illustrate pore lining and strong presence of sodium ions in the pore lumen. (**E**) The inner and outer β-barrel in the cap region (upper arrow in (D)) are held together mainly by hydrophobic interactions. Some key residues are shown. In the center, ring of six E115 residues strongly attracts cations (spheres). Rings formed by six E80 residues (**F**) slightly above membrane plane (middle arrow in (D)) and by six E101 residues (lower arrow in (D)) within the membrane plane (**G**) also strongly attract cations.

**Figure 8:**
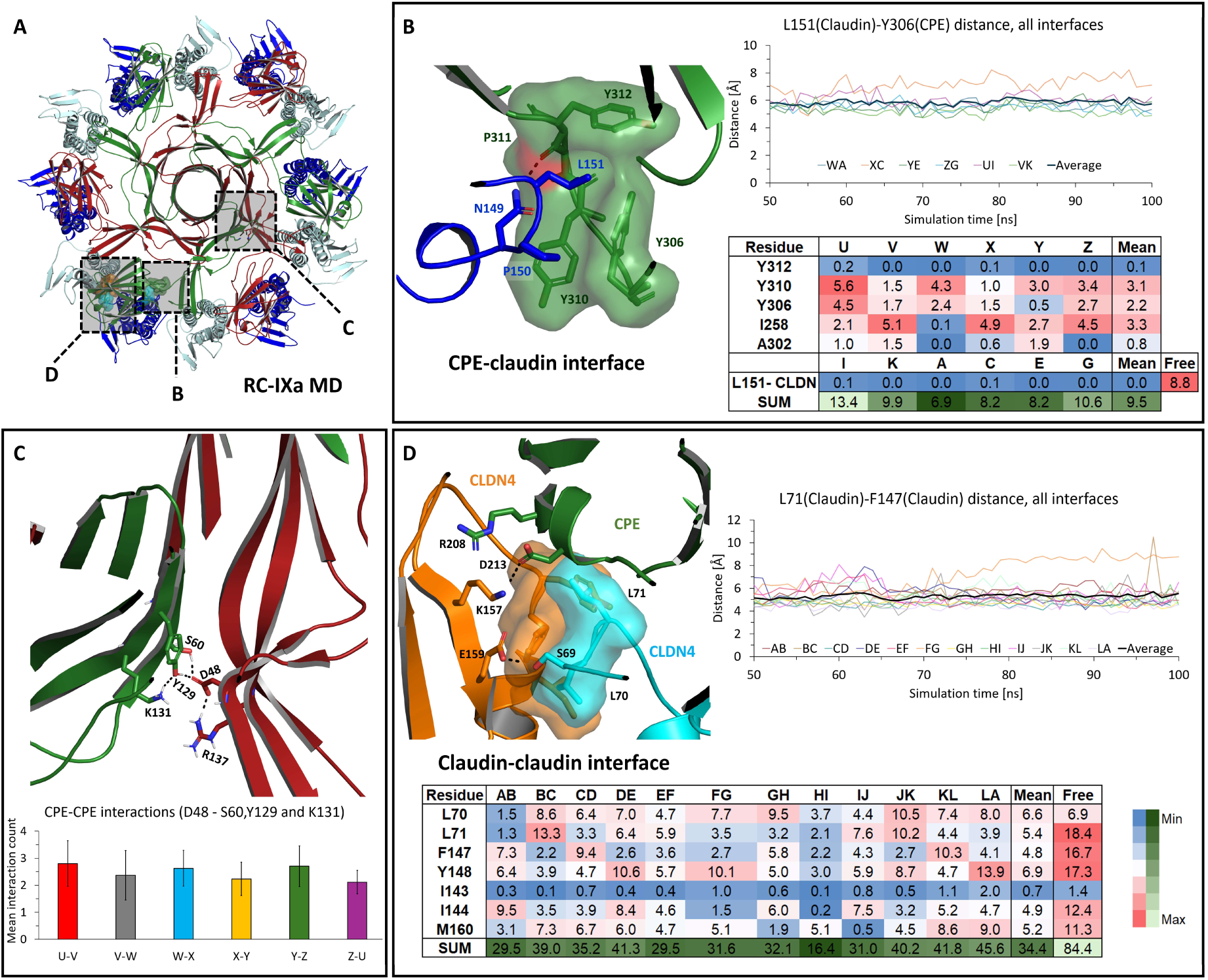
Analysis of inter-subunit interfaces after MD simulation of pore complex state RC-IXa. Structure images show snapshots after 100 ns. (**A**) Top view of complex with CPE (green and red), canonical receptor CLDN4 (blue) and additional CLDN4 (cyan) subunits. Close-up regions shown in (B, C, D) are highlighted. (**B**) Key CPE-CLDN4 interface. Left: Cartoon of interface, key residues are shown as sticks and binding pocket in CPE as surface. Perspective is turned by ∼180° with respect to (A). Top: Distance between L151 (Cγ) and Y306 (Cγ) over last 50 ns of simulation for individual CPE-CLDN4 dimers (CDLN4 chains A, C, E, G, I, K; CPE chains U, V, W, X, Y, Z) and average. Bottom: SASA values (mean for last 50 ns) for the individual subunits (letters) and mean. Free: Mean value of L151 of the six claudin not interacting with this pocket in cCPE. (**C**) CPE-CPE interactions in cap region of pore. Top: Cartoon of interface, key residues are shown as sticks. Bottom: Mean interaction count (<u>±</u>SD) for D48 with S60, Y129 and K131 within the last 50 ns of simulation for the six CPE-CPE subunit interfaces. (**D**) Cis-interface between two CLDN4 subunits. Top left: Cartoon of interface, key residues are shown as sticks and interacting hydrophobic residues also as surface. CLDN4 subunits in orange and cyan, CPE in green. Top right: Distance between L71 (Cγ) and F147 (Cγ) over last 50 ns of simulation for individual CLDN4 dimers (subunits given as letters) and average. Bottom: SASA values (mean for last 50 ns) for the individual subunits (letters) and mean. Free: Mean value of corresponding residues in RC-VIIIa simulation not consisting of CLDN4-CLDN4 interfaces.

**Figure 9:**
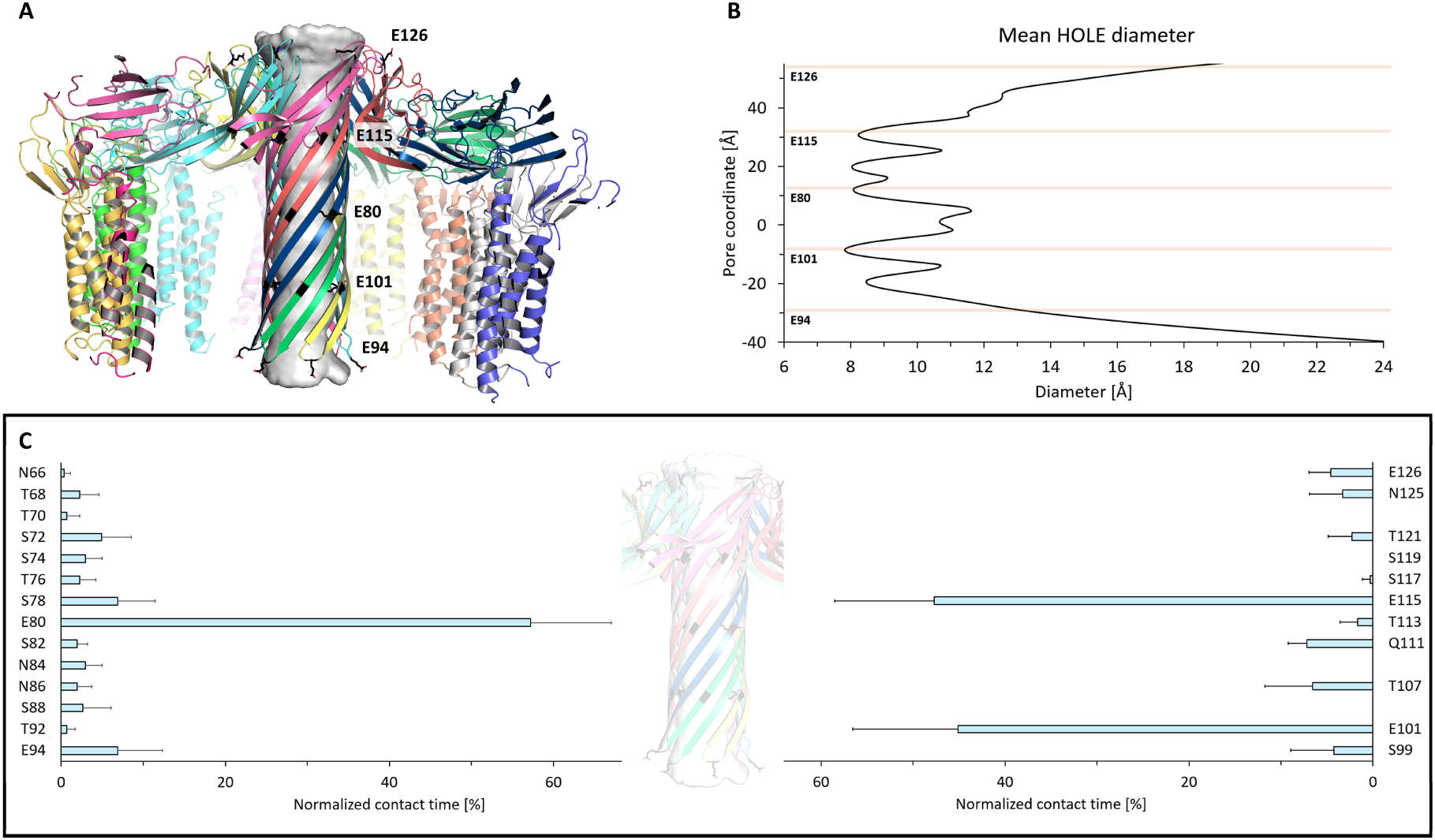
Pore diameter and interaction of pore-lining residues with sodium ions. Quantitative analysis of MD simulation for state RC-IXa. **(A)** Pore path and dimensions (gray surface) of the pore β-barrel, as obtained by HOLE ^66^(snapshot at 100 ns, CPE and Cldn4 subunits shown as colored cartoons, clipped. The positions of pore-lining glutamates are represented in black sticks and cartoon. **(B)** Diameter along the pore coordinates calculated by HOLE. Narrowing of the pore near the hexa-glutamate rings by E80, E101 and E115 could be observed. Mean minimal pore diameter at E101: 7.8 Å. **(C)** Normalized contact times of pore-lining residues with Na^+^ ions over simulation time (last 50 ns) is given. Hexa-glutamates that narrows the pore (E80, E101, E115) interacted more (>40% of time) than the other neighboring polar residues (∼5-10%), suggesting key role of the glutamates in ion permeation. E94 and E126 residing at the pore periphery interacted only ∼5-10% of time, indicating role in attraction of cations towards the pore.

The central stem pore β-barrel (89 Å long) showed a stable conformation (Figure 7A-C, Figure S8) with extended H-bonds between the twelve β-strands throughout the 100 ns simulation run. The β-hairpin tips, including E94, consistently reached towards the cytoplasmic side. Hydrophobic residues of the lipid-facing side of the β-barrel transmembrane region (80-108) remained well-embedded in the membrane core (**Figure 7C, D**). The pore lumen was lined with hydrophilic non-charged and negatively charged residues, filled with water and sodium ions, but hardly with chloride ions (**Figure 7D-G**). Along the pore, three hexa-glutamate rings were formed by E101, E80, E115, respectively. These three rings interacted strongly with Na^+^ ions and each glutamate was surrounded by polar residues, even though the rings differed in the content of glutamine, asparagine, serine and threonine residues (**Figure 7D-G**). The pronounced strong cation attraction fitted well with the experimentally demonstrated high cation selectivity of the CPE pore.^51, 52^

The cap region was stabilized by the two concentric β-barrels (**Figure 7E**). Here, intramolecular contacts between two β-sheets were primarily maintained by hydrophobic interactions (between V58, L63, F71, V118, T120, V128, Y130, V132, I161, L163), as in the monomeric CPE. Intermolecular interactions between CPEs and CPE and CLDN4 were facilitated by H-bonds between neighboring β-strands and by electrostatic side chain interactions (**Figure 8A**). The latter include the E67-K165 interaction located at the top between the inner and the outer β-barrel and at the bottom outer backside, an electrostatic network centered around D48 with residues including S60, Y129, K131, R137, T159 (**Figure 8C**). Importantly, this fitted perfectly to experimental data which show that D48 is essential for CPE pre- pore formation.^65^ Thus, this interface was quantified in more detail by measuring the mean electrostatic interactions between D48 of one subunit and S60, Y129 and K131 of a neighboring subunit for all CPE subunit within the last 50 ns of the simulation (**Figure 8C**). The mean interaction count for the six CPE subunit pairs was ∼2.47±0.25, suggesting a significant contribution to the CPE oligomer stabilization.

Concerning the 12 claudin subunits, their transmembrane four helix bundles remained also well-embedded in the membrane throughout the simulation. Thus, importantly, in contrast to the claudin-anchored state RC-VII (**Figure 3A**) the claudins as well as the CPE pore were spanning the same membrane plane.

In addition, the interaction between cCPE and CLDN4 was highly stable, including the interfaces observed in the experimental cCPE/CLDN4 structures (PDB 3X29, 5B2G, 6AKG, 6AKE, 6OV3, 6OV2, 7KP4, 8U4V; ^33–35, 41–43^. In particular, L151 in the ECS2 of CLDN4 was stably plugged into the triple tyrosine pocket of cCPE (**Figure 8B**, ^32, 33^). This claudin ECS2-cCPE interface was analyzed quantitatively. The distance between the residues L151 (Cγ, CLDN4) and Y306 (Cγ, CPE) were measured over the last 50 ns of the simulation time (**Figure 8B**, top right). The values remained nearly constant over time, and except for one interface, all the interfaces had distances between 5 and 6 Å, with a mean distance for all six CLDN4-cCPE interfaces of 5.85±0.61 Å. To quantify the hydrophobic interactions in this region, the solvent accessible surface area (SASA) of residues involved in such interactions (I258, A302, Y306, Y310, and Y312 from CPE, and L151 from CLDN4) were calculated (Figure 8B, lower right). SASA values were normalized with respect to the number of C atoms in each residue. The exceptionally low values suggested significant water exclusion in the region across all interfaces. Notably, the SASA of L151 from CLDN4 which is embedded into the cCPE pocket, was 0.0 – 0.1, indicating total water exclusion. Similarly, the SASA of residue Y312 from CPE, which is present well inside the hydrophobic pocket, was nearly as low. In total, the distance and SASA values suggest that these residues form a highly hydrophobic interface that is stably maintained over time.

Furthermore, the claudin-claudin interaction within the dodecamer ring was analyzed. Despite variations observed between the different subunit-subunit interfaces and over time, a constant association between the subunits was observed. Similar to the linear cis-interaction in claudin strands ^55, 56^, proximity of hydrophobic residues as well as electrostatic interaction were observed between residues of ECS2 (E159) and ECH region (S69) **(Figure 8D**). For the linear-cis-like interaction, the distance between the residues L71 (Cγ) and F147 (Cγ) from two neighboring claudin subunits was calculated across all 12 interfaces (**Figure 8D**). These distances remained stable among the interfaces over the last 50 ns of the simulation time, except for one interface which showed a slight increase in distance from ∼6 to ∼8 Å. Overall, the mean distance between L71-F147 across the interfaces was ∼5.29±0.17 Å, suggesting stable cis interfaces. This was further supported by the SASA values of residues that take part in the cis interactions, which were comparable to similar SASA calculations of linear-cis interfaces in the claudin strand models.^55, 56^ For L71 from ECH of a claudin subunit, the residue that fits inside the hydrophobic pocket of the cis-interacting claudin, a low SASA value was calculated. The neighboring L70 from ECH had a slightly higher SASA (mean values of 5.4 and 6.6, respectively). In addition to L70, other residues involved in hydrophobic interactions in the interface like F147 and M160 had slightly higher SASA than their counterparts in the strand models. Overall, the distance and SASA measures indicate stable yet alternative linear-cis interfaces in the claudin ring of CPE-claudin complex, which are partly comparable to such interactions observed in the claudin strand models. Of interest, frequently cCPE residues also appeared to stabilize these claudin-claudin interface through interactions between cCPE-D213 and CLDN4-ECS2-K157, and cCPE-R208 with the CLDN4-ECS2 turn region (Figure 8D).

### Detailed analysis of the ion permeation pathway of the CPE pore

The ion permeation pathway of the CPE pore was further analyzed using the state RC-IXa simulation trajectory data and the HOLE program (**Figure 9**). On analyzing the mean pore diameter profile (**Figure 9B**), we observed narrowing of the pore near the three hexa-glutamate rings formed by E80, E101, and E115, respectively. Particularly, the minimal constriction zone was observed adjacent to the E101 ring that is closest to the cytoplasmic side with a mean diameter of 7.8 Å (**Figure 9B**).

To investigate the interaction of pore-lining residues with incoming ions, we calculated the normalized contact time of these residues with Na^+^ ions over time (**Figure 9C**). The glutamate residues E80, E101, and E115 exhibited prolonged contact times, each interacting with Na+ ions for more than 40% of the simulation duration. Additionally, neighboring polar residues on either side of these strongly interacting glutamates, showed shorter contact times (∼5-10%). This suggests that these residues likely play supporting roles in ion transport. Of note, these residues differ between the three glutamate rings. In contrast to E80, E101, and E115 residing within the pore, E94 and E126 located at the periphery of the pore, showed short contact times (∼5-10%). These residues might contribute to cation attraction towards the pore barrel.

In total, the MD simulations strongly supported the modeled architecture, claudin anchorage, membrane embedment and pore lining of the CPE pore, as well as a key role for E80, E101 and E115 residues in the cation selective ion permeation through CPE pores.

### Cellular binding and viability assays with (c)CPE-incubated cells indicate interaction between claudin subunits during assembly of CPE pore complex

The models, in particular state RC-IXa, indicated an interaction between CLDN4 subunits similar to the highly conserved linear cis-interaction in polymeric claudin TJ strands.^14, 55, 56, 67^ Previously, we showed that glutamate mutation of S68 in CLDN3 (corresponding to S69 in CLDN4) at the conserved linear cis-interface inhibits claudin oligomerization. ^68^ Given, the high sequence similarity between CLDN3 and CLDN4 (89% without C-terminal cytoplasmic tail) and the finding that CPE binds strongly to both claudins^32^, we took advantage of the existence of the previously characterized HEK293 cell lines expressing either CLDN3wt or CLDN3S68E but no endogenous CPE-binding claudins^32, 68^, and established cellular binding and viability assays.^15, 32^ These were used to test whether weakening claudin cis-interaction affects binding of cCPE to claudins and/or pore complex formation by CPE. Functional cytotoxic pore formation has a direct effect on cell viability, measured by the cells’ ability to metabolize MTT. Both cell lines, Hek-CLDN3WT and Hek-CLDN3-S68E, previously treated with different concentrations of CPE, were susceptible to a decrease in viability with increasing CPE concentration (**Figure 10A**). However, S68E mutation slightly inhibited this CPE-mediated cell damage of CLDN3-expressing cells, which is visible from higher EC_50_ values (0,52±05 nM forCLDN3WT and 1,24±0,11 nM for CLDN3-S68E, **Figure 10A,B**). In binding saturation experiments (**Figure 10C**, representative experiment out of 3), the concentration-dependent binding of cCPE to HEK-293 cells expressing CLDN3 or CLDN3-S68E was measured and used to determine K_D_-values as the cCPE concentration at half-maximal binding (**Figure 10C, D**). In addition to its weakening effect on CPE-mediated cell damage, the S68E mutant slightly decreased the affinity of cCPE for CLDN3 (K_D_: 14,09±5,07 nM for CLDN3WT and 37,66±3,33 nM for CLDN3-S68E, **Figure 10D**). These in vitro data support the idea that an interaction related to the linear-cis interaction in claudin strands contributes to the formation of claudin-anchored CPE-pore complexes.

**Figure 10:**
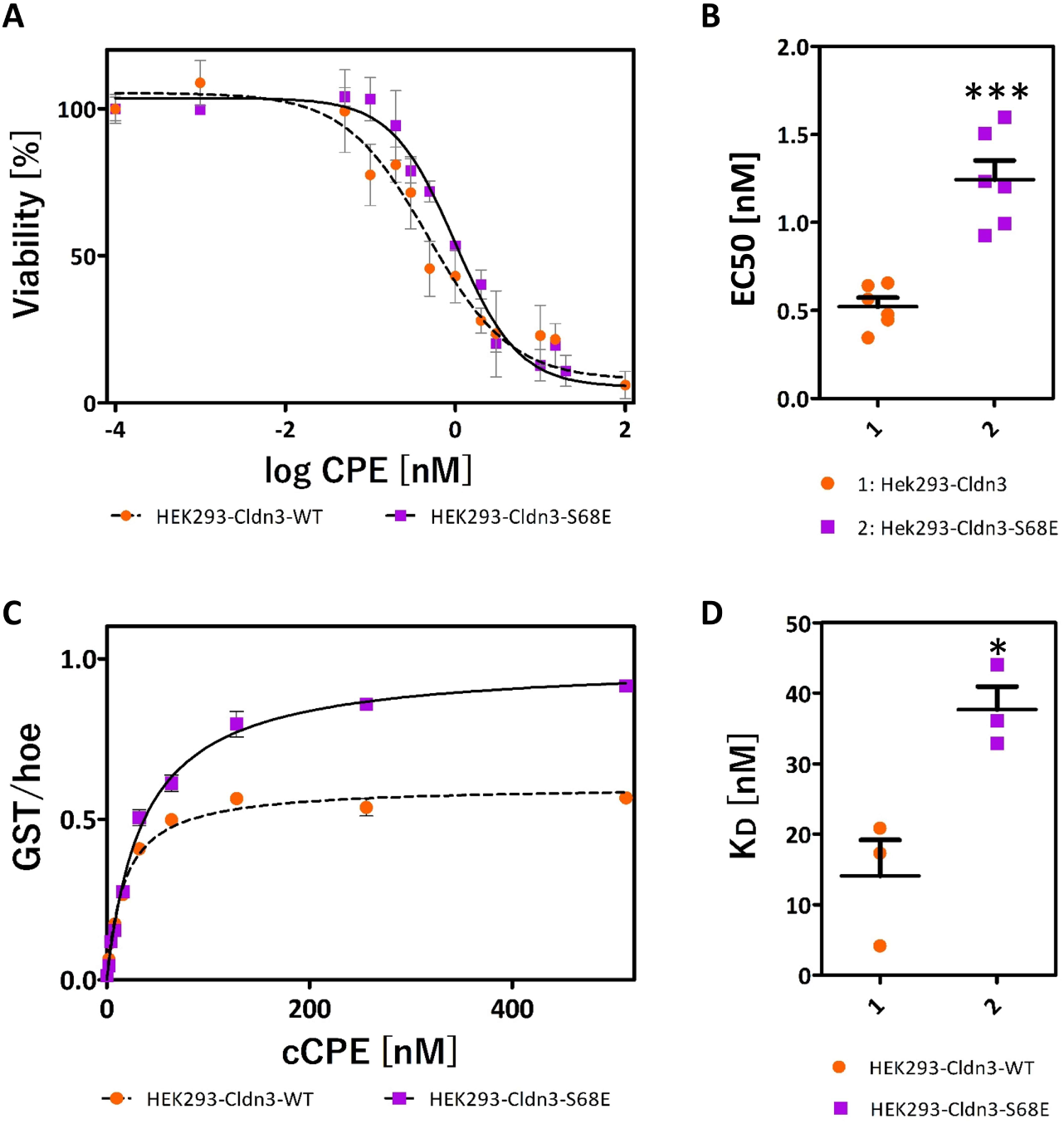
S68E mutation in CLDN3 at conserved linear cis-interface weakens CPE-mediated cell damage **(A, B)** and binding of GST-cCPE **(C, D)** compared to CLDN3-WT in stably transfected HEK293 cells. **(A)** Viability of CLDN3-WT- or CLDN3-S68E-expressing HEK cells after 1 h incubation with different CPE concentrations tested by MTT assay. Normalized values (n=3) were plotted against log of concentration (nM). **(B)** EC50 calculation revealed a significantly higher EC50 value for HEK293 cells expressing the CLDN3-S68E mutant. Mean±SEM, n≥6, *** p < 0.0003. **(C, D)** Cellular binding assay. CLDN3-WT- or CLDN3-S68E-expressing HEK293 cells, or non-transfected, claudin-free HEK293 cells (nt) were incubated for 30 min with different concentrations of GST-cCPE, after fixation, binding was detected by anti-GST antibodies and normalized to cell number (Hoechst signal). From the binding data (double determination), the unspecific binding (binding to nt cells) was subtracted. The K_d_ was significantly higher for the CLDN3-S68E mutant. Mean±SEM, n ≥3, * p < 0.05.

## Discussion

In this study, we predicted atomic and derived mechanistic models of different states of oligomerization and prepore-to-pore transition of the cytotoxic CPE-claudin pore complex. Alignment of different predicted structures obtained with an AlphaFold2 oligomer prediction platform and experimentally solved CPE and cCPE/claudin structures was combined with published biochemical data on the CPE-claudin pore complex and MD simulations to drive model building.

The key features of the developed model for assembly of the CPE-pore-complex are schematically summarized in **Figure 11**. (1.) CPE monomers bind to claudin monomers or dimers containing one or two high-affinity (e.g. CLDN4) and one or none low-affinity receptors (e.g. CLDN1). A trimeric complex is stabilized by (a) the established high-affinity Claudin-cCPE interaction ^32–34^, (b) an experimentally indicated CLDN-CLDN interaction (modified linear-cis-interaction, Figure 10), and a proposed additional low-affinity CLDN-cCPE interaction (**Figure 8D**, **Figure 11B, F**). This trimer is proposed to represent the small complex identified by co-immunoprecipitation analysis.^38, 39, 60^ (2.) With only minor conformational changes 6 of these small complexes oligomerize into a large pre-pore complex containing 12 claudin and 6 CPE subunits **(Figure11 C, G**, states RC-III/RC-IV ). This stoichiometry was suggested by heteromer gel shift, size exclusion chromatography and co-immunoprecipitation analysis.^39^ (3.) The α1-helix swaps clockwise from domain II in the same CPE subunit to domain II in the neighboring subunit. Concomitant anti-clockwise movement of β-sheet in CPE domain II but not of the claudin receptors support change of angle between domain II and cCPE (**Figure 11 D, H**). (4.) The α1-helix dissociates from the domain II β-sheet leading to stepwise twisted extension of the inner β-barrel of CPE (state RC-VI to RC-IX, (**Figure 11 I-N**). (5.) Further conformational changes are required to form and drive the tips of the extending β-barrel through the membrane. nCPE has to get closer to the membrane without disturbing the cCPE-receptor (claudin) interaction. The model shows that this can be obtained by a conformational change of the flexible linker between cCPE (domain I) and nCPE (domains II&III) (**Figure 11L, M, N**). (6.) The diameter of the ring of 12 claudins has to be reduced to form the proposed continuous dodecameric claudin ring. The models show that this can be obtained by an additional conformational change in the cCPE-nCPE linker region resulting in a kink between cCPE and nCPE (**Figure 11D, I, J**), and by a variant of the claudin-claudin cis-interaction (ECH-ECS2 pocket) suggested to drive polymerization of claudins within TJ strands.^14, 55, 56^ We propose that the mechanistic advantage of such a receptor ring is that it may support a twisting-like rotational movement of the central nCPE β-barrel against the interjacent CPE domain II and the peripheral cCPE-claudin ring of the complex. This might drive β-barrel extension and penetration through the membrane (**Figure 11 D, H-N**). Thus, our structural model provides a mechanistic model of the CPE-claudin pore formation to be further tested and validated by future studies.

**Figure 11:**
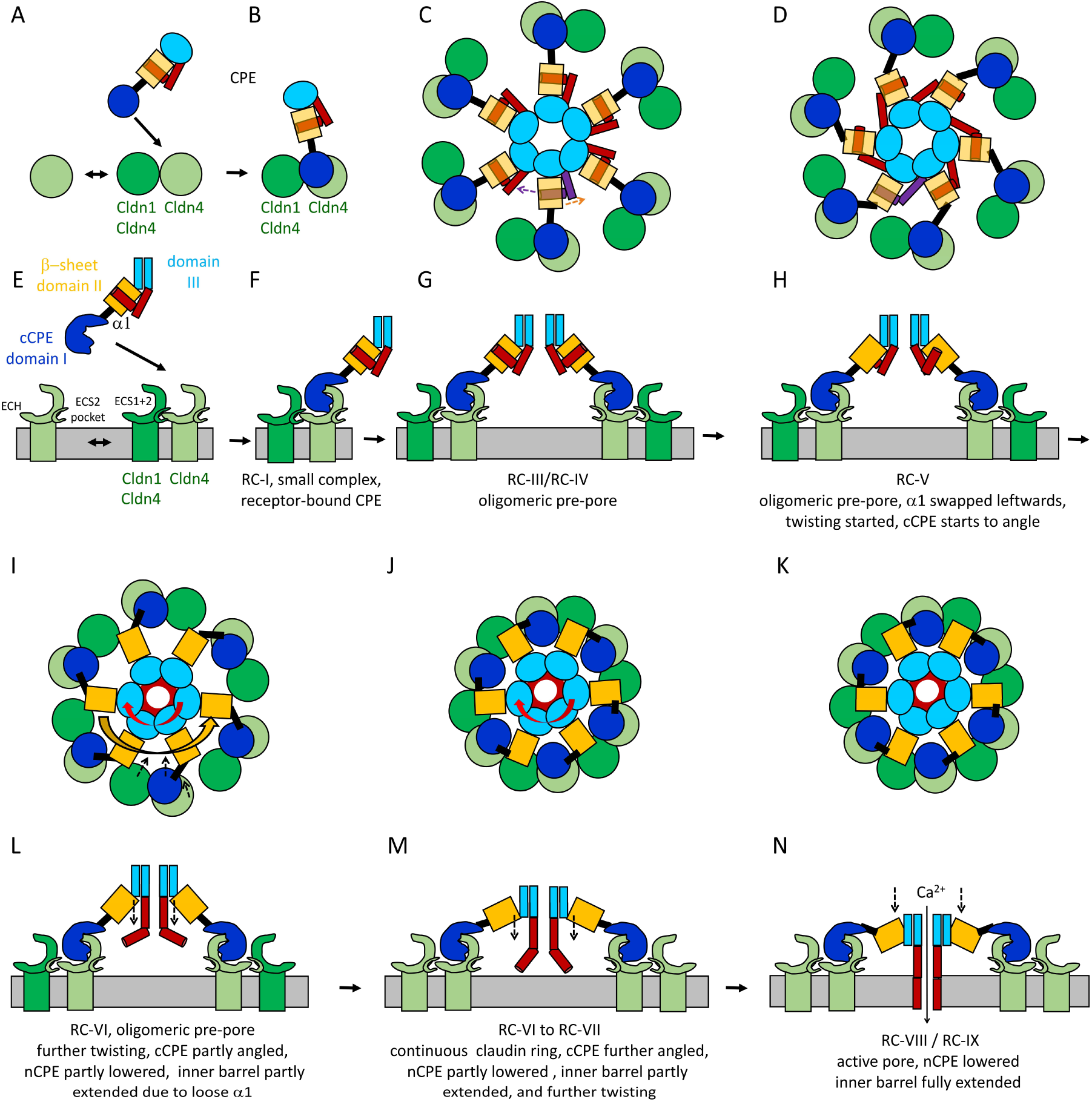
Schematic model of CPE-claudin pore complex assembly. Top views (A, B, C, D, I, J, K) and respective side views (E, F, G, H, L, M, N) of different stages. **(A, E)** Claudin dimers form dynamically. **(B, F)** Monomeric CPE binds to claudin dimer containing one or two high affinity (e.g. Cldn4) and one or none low affinity receptor (e.g. Cldn1). The trimeric complex is stabilized by high affinity Cldn4-cCPE-, Cldn-Cldn cis-, and low affinity Cldn-cCPE interactions (small complex). **(C, G)** 6 small complexes oligomerize with minor conformational changes, only, into a large pre-pore complex (CH-I) containing 12 claudin and 6 CPE subunits. (**D, H**) α1-helix swaps clockwise from domain II in same CPE subunit to domain II in neighboring subunit. Concomitant anti-clockwise movement of β-sheet in domain II, but not of the claudin receptors, support change of angle between domain II and cCPE. (**I, L**) α1-helix dissociates from the domain II β-sheet leading to twisted extension of inner β-barrel. nCPE (domains II&III) starts to move downwards to membrane. cCPE angles further relative to nCPE. Claudins bound to cCPE move more to center. (**J, M**) cCPE angles and moves further relative to nCPE leading to formation of a continuous claudin ring. (**K, N**) Further twisting movement of complex center against complex periphery drives full extension of the β-barrel, membrane penetration and formation of the active pore.

The different components of the model are discussed below.

### Formation of a hexameric CPE β-barrel pore

The model of the hexameric nCPE pre-pore state (state nCPE-IV, **Figure 1F**) shows very high similarity to the HA3 subcomponent of *C. botulinum* Type C Progenitor Toxin hetero-oligomer (**Figure 2D, E**). This state is highly reliable, due to the highly similar fold of both proteins (RMSD for protein backbone of CPE26-202 and HA3a or HA3b: 1.79 Å), similar oligomeric arrangement (RMSD for protein backbone of nCPE-IV (CPE26-202) and HA3a oligomer: 3.14 Å) and formation of a similar concentric double-β−barrel as found in other aerolysin-like toxins.^46–48, 69^ In addition, the hexameric active pore states (nCPE-VII, RC-VIIIa and RC-IXa) show a similar elongated β-barrel pore arrangement as other aerolysin-like toxins (^46–48, 69^, **Figure 2**). However, in contrast to aerolysin, the unique barrel consists of 12 β-strands originating form 6 CPE subunits instead of 14 β-strands originating form 7 toxins subunits. Importantly, the hexameric pore was stable during multiple MD simulations (for nCPE-VII, RC-VIIIa, RC-VIIIb, RC-IXa, and RC-IXb, **Figure 4**, **7**, **8**). Compared to the corresponding heptameric CPE pore model, the hexameric one was more favorable (**Figure S2**) and were much better in agreement with the biochemical data that indicate a hexamer ^39^.

The reliability of the modeled pore β-barrel is also supported by the following: (i) The overall mushroom-like architecture consisting of an elongated β-barrel stem (∼9 nm long and ∼ 3.5 nm wide) and a cap with a central concentric double β-barrel and receptor binding domains at the periphery is similar to other aerolysin-like toxins.^26, 47, 48^ (ii) The hydrophobicity of all membrane-facing and hydrophilicity of all pore lumen-lining residues. (iii) Complete consistency to experimental data: (a) Residues 80-106 form the membrane-spanning β-hairpin region with F91 and F96 at the tip (**Figure 7**), as shown by site-directed mutagenesis (region 80-106, residues F91, F95, G103 are critical for membrane insertion^40, 70^; (b) D48 participates directly in CPE-CPE interface (**Figure 8C**), fitting to its essential contribution to oligomerization^11^; (c) three unique hexa-glutamate rings along the ∼8 Å wide pore are formed by E101, E80 and E115 which strongly attract cations leading to charge-selectivity fitting to the strong cation selectivity of the CPE pore demonstrated by electrophysiological measurements.^51, 52^

In contrast to other members of the aerolysin family (like aerolysin, lysenin, or *C. perfringens* ɛ-toxin), the membrane-spanning region of CPE needs to undergo an α-to-β transition of secondary structure during pore formation. Thereby, the α-helix of domain II changes its conformation. Similar changes have been shown for β-barrel pore formation in cholesterol-dependent cytolysins, driven by π-stacking and electrostatic interactions, and was so far described as a unique feature of the CDC/MACPF/SNTX (cholesterol-dependent cytolysin/membrane attack complex perforin/stonefish toxin) superfamily of PFTs.^71^ The opposite, a β-to-α-transition of the pore forming region, occurs in the α-PFT Cytolysin A (ClyA).^72^

Compared to the nCPE structures in the states nCPE-IV, nCPE-VII, RC-VIIIa, and RC-IXa, the structures in the intermediate states nCPE-V and nCPE-VI (Figure 1) are based on less corroborating data. However, the combination of AlphaFold2 predictions with known structures has enabled us to deduce a potential sequence of conformational changes in nCPE, from the monomeric CPE to the active transmembrane pore complex.

We also tested Alphafold3 with the same CPE inputs as used for Alphafold2 (sequences, number of subunits, see **Table 1).** No meaningful hexamers or heptamers were obtained that contained a central prepore or pore formed by nCPE. Strikingly, addition of CLDN4 to CPE, either in a 1:1 or 2:1 stoichiometry, yielded prepore-like complexes with cCPE/Cldn4 interface very similar to the one in dimer crystal structures.^34, 57^ However, in contrast to the Alphafold2 models (nCPE state IV to VII) they did not contain a β-barrel (e.g. formed by β3- and β5 strands of nCPE). Moreover, D48 was not part of an oligomeric interface and the resulting tilt of the claudins in the membrane was very different from the one in claudin strand models^55, 56, 63^ (see **Figure S9** for an example). Thus, the reliability for the Alphafold3 prepore models is questionable. Interestingly, a very recent study^73^ also used Alphafold3 to generate very similar CPE prepore models. The main focus of this work is the structure of the dimeric CPE-CLDN4 complex that was revealed by cryo-EM. In addition, the authors propose that the tilted claudins in the Alphafold3-predicted prepore complex could cause a positive membrane curvature that could support insertion of the membrane-spanning CPE segment during pore formation. Such a membrane curvature could also facilitate that the β-hairpins in our pore model state RC-VII are fully spanning the membrane (**Figure** 3A).

**Table 1:**
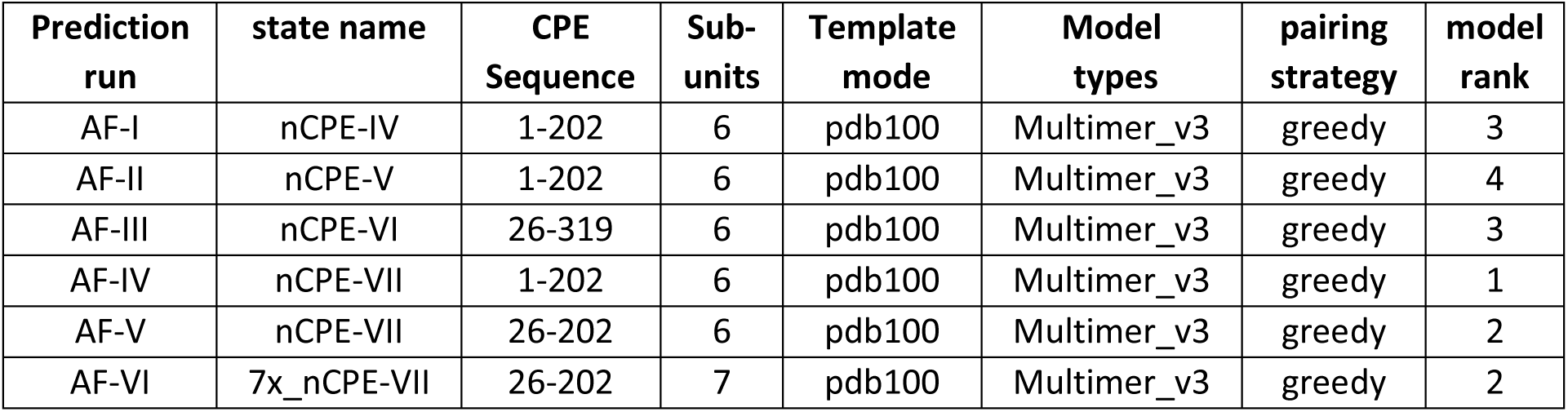
AlphaFold (AF) prediction runs.

### CPE-claudin interaction

The anchorage of the CPE-hexamer to claudin receptors in the membrane fits very well with respect to topology relative to the membrane and to the known interaction sites between cCPE and claudins. In particular, the experimental biochemical and structural data identified key interactions (CLDN4-ECS2-P150/L151 within Y306/Y310/Y12 pocket in cCPE and CLDN4-ECS2-N149 with cCPE P311 backbone) that remained very stable throughout MD simulations of several membrane embedded oligomeric CPE-CLDN4 complexes (RC-VIIIa, RC-VIIIb, RC-IXb and for RC-IXa shown in **Figure 8B**). Thus, this CPE-claudin interaction is of high reliability. Of note, to our knowledge, this is the first time that stability of the CPE-claudin interaction was demonstrated on the atomic level under dynamic conditions. Interestingly, in state RC-IXa containing the putative dodecameric CLDN4 ring, an additional interaction site was found between CPE and a second, neighboring claudin (cCPE-D213 with CLDN4-ECS2-K157 and cCPE-R208 with the CLDN4-ECS2 turn), leading to a trimeric CPE-CLDN4-CLDN4 interaction (**Figure 8D**, **11B**). Importantly, the additional CPE-claudin interaction would not depend on the ECS2 interaction motif that is found only in canonical CPE receptor claudins, such as CLDN3, -4, -6 to -9,-14.^16, 37, 74^ Thus, also CLDN1 could be a co-receptor, explaining why CLDN1 is robustly identified in CPE-claudin complexes despite its low affinity to cCPE.^38, 39, 74, 75^ However, the validity of the proposed additional CPE-claudin interface as well as of the proposed claudin-claudin cis-interface for the small complex and the active pore complex remain uncertain, because also other subunit arrangements are thinkable, for instance with face-to-face claudin dimers^14^ that would be hardly compatible with formation of a continuous claudin ring.

### Conformational changes in the nCPE-cCPE linker region during CPE-claudin pore complex assembly

The nCPE pore predictions and simulations as well as the published cCPE-claudin structural interaction data appeared to be very reliable. However, combining both with the nCPE-cCPE linkage present in the crystal structure of monomeric CPE did not result in a complex in which the pore can fully span the membrane plane that is defined by the transmembrane domains of the claudins (state RC-VII, **Figure 3A**). Of note the β-strands -1, -2, -6, -7 that connect domain II and III in nCPE did not change strongly between the monomeric CPE and state nCPE-VII (and also RC-VII to RC-IX) (**Figure S1, 1, 3, 6H**). Thus, the most plausible conformational change to enable a transmembrane pore appeared to be a modification of the probably more flexible nCPE-cCPE linker region. Consequently, in a first step, nCPE was move mainly downwards relative to the cCPE/CLDN4 complex (state RC-VIII, **Figure 3**) which resulted in a complex with a stable and functional transmembrane pore (F**igure 4F-M**). In a second step, cCPE was further re-arranged relative to nCPE in order to fit the 6 CPE subunits to a continuous ring of 12 claudin subunits (state RC-IXa, **Figure 5**, **6**, **7**) that is here proposed to provide mechanical support (**Figure 11**). Although these manual modifications extensively changed the overall CPE conformation, they did neither alter those of the hexameric nCPE nor those of the cCPE/CLDN4 complexes. In addition, in the CPE-related HA3 complex (F**igure 2E**), the domain corresponding to cCPE is also angled slightly in the same direction. Due to the manual modification of the nCPE-cCPE linker region, details in the conformation of the linker and in the relative positioning of cCPE, nCPE and the claudin not bound to the Triple-Tyr pocket of cCPE are more uncertain than other aspects of the complex. Nevertheless, the fact that the resulting structure (state RC-IXa) led to stable and functional receptor-bound cation-selective oligomeric CPE pores in MD simulations provides at least a plausible model for conformational changes in the nCPE-cCPE linker during pore-formation.

### Driving force for transmembrane pore barrel formation

We propose that formation of a receptor ring may support a twisting-like rotational movement of the inner nCPE pore β-barrel against the outer cCPE-claudin ring. This could contribute to β-barrel extension and/or its penetration through the membrane. Further analysis, particularly through comparisons with the pore formation mechanisms of other β-PFTs, is expected to validate and elucidate the details of this mechanism. The postulated overall CPE prepore-to-pore conversion resembles that of other aerolysin-like β-PFTs: Prepore formation, β-barrel expansion in a zipper-like manner and collapse by change of angle between the receptor-binding and the pore-forming domains.^47, 76^ Interestingly, much like in our CPE model conformational movements including rotation of central against peripheral domains and flattening of the complex have been observed for *C. perfringens* ε−toxin^77^ and aerolysin.^78^ In aerolysin, rotation of domain 4 potentially drives the circular association of β-sandwich domains of each monomer, leading to (prepore-) β-barrel formation. This driving force hypothesis was derived through structure determination of different states (prepore to final pore), analogous to our CPE study in which different states were modeled.^26, 78^ In addition, alternating polar serine and threonine residues in the insertion loop of all β-PFTs may contribute to driving the prepore-to-pore transition by aiding membrane binding, oligomerization, and facilitating amphipathic loop insertion during transmembrane pore formation.^47^ Further studies could take up a more in-depth, comparative analysis regarding this transition using our CPE model, which unlike other studies on PFTs, also strongly considers the toxin receptor proteins.

### Diameter and selectivity of the CPE pore

When analyzing the mean pore diameter profile (**Figure 10B**), we observed the minimal constriction zone adjacent to the E101 ring closes to the cytoplasmic side with a mean diameter of 7.8 Å (**Figure 9B**). Benz and Popoff estimated that the diameter of a CPE channel could roughly be 14 Å, wider than in our model.^52^ They determined this value through a fit of the single channel conductance as a function of the bulk solution. However, such a discrepancy in experimentally calculated pore diameters with the actual diameters in the resolved structures was already seen in the case of other β-PFTs.^79^ For example, different studies reported varying diameters from 7 Å to 30 Å for the aerolysin pore (30 Å ^80^, 7 Å ^81^ ; 9.3 Å ^82^; ∼17 Å ^83^; 17 Å ^84^; 19-23 nm ^85^), but the actual diameter observed in the aerolysin pore crystal structure (PDB ID: 5JZT) was 13.7 Å.^47^ In addition, Benz and Popoff showed that the CPE pores are highly cation selective, whereas most other β-PFTs are either anion-selective or non-selective.^52^ This unique cation selectivity was clearly highlighted in our model.

## Conclusion

In summary, we predicted a model of CPE pore formation that is fully consistent with published structural, biochemical and functional data. However, limitations of this study are that it does not contain detailed experimental validation by *in vitro* structure-function analysis of CPE and claudin mutants nor a new experimentally solved complex structure. Nevertheless, it provides for the first time a detailed structural and mechanistic model for the pore formation and charge selectivity of the CPE-claudin pore complex. This novel knowledge opens the way for further investigations on the structure, physiology and pathophysiology of CPE and claudins.

## Materials and Methods

### Modeling and MD simulation of CPE-CLDN4 complexes

The Schrödinger Maestro BioLuminate software (BioLuminate, version 4.9.132, Release 2022-4, Schrödinger, LLC, Germany, 2022) was used to model, refine, and simulate the CPE pore – CLDN4 models. All the simulations were conducted on a Linux-x86_64-based GPU workstation. Both BioLuminate and Schrödinger PyMOL 2.5.7 (http://www.pymol.org/) were utilized to create the model representations presented in the figures.

### AlphaFold2-based structural prediction of CPE oligomers

For generation of initial models of the oligomeric CPE pore, the AlphaFold2 and MMseqs2 based prediction platform ColabFold v1.5.5 (https://colab.research.google.com/github/sokrypton/ColabFold/blob/main/AlphaFold2.ipynb) was used with GPUs, extended RAM and Python 3.^49^ The following input parameters were varied: (a) Sequence: 1-319, 1-202, 26-202, 26-319; (b) number of subunits: 6 or 7; (c) template mode: none, pdb 100, custom; (d) model types: Alphafold2_multimer_v3, Alphafold_ptm, deepfold_v1; (e) pairing strategy: greedy, complete; (f). For other parameters, the default settings were used. The following AlphaFold (AF) prediction runs were used for further analysis:

### Further modeling, multi subunit assembly and MD simulation of CPE and CPE-CLDN4 complexes

From the structure AF-IV (= state nCPE-VII), the disordered N-terminal region (M1-K25) was removed, the structure was prepared, and embedded in a membrane lipid bilayer made up of 1-palmitoyl-2-oleoyl-sn-glycero-3-phosphocholine (POPC) lipids. The positioning of the CPE pores within the membrane relative to the hydrocarbon layer of the lipid bilayer was calculated using the PPM 3.0 web server^54^ (https://opm.phar.umich.edu/ppm_server). The membrane-protein complex was placed in an orthorhombic simulation box with buffer distances of 10 Å, 10 Å, and 15 Å in x-, y-, and z-axes, respectively. TIP3P water molecules^86^, charge neutralizing Na^+^ ions, and 0.15 M Na^+^Cl^-^ salt were added to generate the simulation system. The simulations were carried out using the Desmond Molecular Dynamics module (Desmond Molecular Dynamics System, D. E. Shaw Research, New York, NY, 2022; Maestro-Desmond Interoperability Tools, Schrödinger, New York, NY, 2022;^87^) within Maestro BioLuminate.

After minimizing the system to a local energy minimum through a 200 ps minimization, it was relaxed in an NPγT ensemble to room temperature, pressure and surface tension – 310 K, 1.01325 bar, 4000 bar. Å. The relaxed system was then equilibrated by stepwise releasing constraints on the protein over ∼45 ns. A final production run, free of constraints, was performed for 100 ns. The simulations were performed using an NPT ensemble in OPLS4 force field^88^, with the Martyna-Tobias-Klein method^89^ and Nosé–Hoover chain method^90^ serving as the barostat and thermostat, respectively.

### Modeling and MD simulations of CPE pore-CLDN4 complexes

#### CPE pore complexes containing six CLDN4 subunits

We developed a CPE pore-CLDN4 complex with six claudin subunits, each bound to the cCPE domain of one of the six CPE subunits. Two variants of CPE pore-CLDN4 complex were modeled, differing mainly in the region between residues V187-A204 in nCPE linking it to cCPE. In one variant (RC-VIIIa), this region forms a β-strand, while in the other (RC-VIIIb), it forms an α-helical conformation.

RC-VIIIb was based on the AlphaFold prediction AF-III (2.1.1.) for CPE_26-319_. However, the β-barrel pore in AF-III was not valid, as it would not span the membrane. Hence, from AF-III, only the residues T26-L63 and Y127-F319 from domain II and III (Figure S1), and cCPE (domain I) were used for modeling the CPE-claudin complex. For the pore, the β-barrel pore structure (N64-E126) from the pore structure simulation (state nCPE-VII after equilibration) was grafted onto AF-II. The crystal structure of human CLDN4 in complex with cCPE (PDB ID: 7KP4) was then aligned with and replaced the cCPE in each CPE subunit of AF-II-nCPE-VII fusion. This modeling resulted in a discontinuous ring of six claudins (receptors) anchoring the full-length CPE hexameric pore (complex RC-VIIIb).

Similarly, another CPE pore-CLDN4 complex (RC-VIIIa) was modeled using the CPE structure AF-IV (=nCPE-VII) instead of AF-III. AF-IV was aligned with RC-VIIIb for identifying the position and subsequent alignment of CLDN4 crystal structures bound with cCPE. One critical difference between RC-VIIIb and RC-VIIIa is that, in RC-VIIIa, the residues (K197-A205) that connect cCPE (from RC-VIIIb) to nCPE (from AF-X1) was manually modified to facilitate the connection.

Each of the two variants (RC-VIIIa and RC-VIIIb) were embedded in a membrane lipid bilayer as described for state nCPE-IV (2.1.). Importantly, the claudin transmembrane helices as well as the CPE pore tips were spanning the membrane in both variants. Following a similar MD simulations protocol as for the nCPE-VII, the two complex systems were minimized, relaxed, equilibrated, and simulated without constraints for 100 ns.

#### CPE pore complexes containing a 12-claudin ring

A second CPE pore-CLDN4 complex was developed with a ring of 12 CLDN4 subunits. However, while modeling such a 12-claudin ring based on the 6-claudin ring (RC-VIIIa), we noticed that the gap between claudins so wide that a contact between neighboring claudins was not possible. Hence, we modeled a 12-claudin ring with a smaller diameter than that of the 6-claudin ring to form a ring with contacts between claudins that represent a modification of the linear-cis interfaces found in claudin strands. For this, our previously published claudin strand model ^55, 56^ was used as a template. A dimer from the template was extracted, and two human CLDN4 structures were aligned with the dimer. The missing N-terminal residues (M1-M4) in the structure from PDB ID 7KP4 was grafted from the PDB ID 5B2G. At first, one of the two subunits in the dimer was rotated by ∼30°. This rotation was necessary to generate an eventual 12-claudin ring by replication. To be more precise, the rotated dimer was duplicated and subunit 1 from the duplicated dimer was aligned with the subunit 2 from the original dimer. Similarly, such rotation and alignment were followed six times. This resulted in a 12-claudin ring connected through an alternative cis interface slightly different than the one found in the strand models. For the orientation of the claudin ring in the membrane plane, we used the membrane from the strand model.^55, 56^ Such modeling also required a more slanted conformation of the regions T26-L63 and Y127-F319 from domain II and III while maintaining the structural integrity of the other CPE elements. In addition, the smaller ring diameter meant insufficient space to maintain the rather elongated CPE conformation. So, each cCPE was further deviated from nCPE and moved from one to the neighboring claudin, and the linker (V187-A204) was modified and connected to nCPE. This resulted in tightly packed nCPE domains with cCPE domains occupying six alternating claudins, while the other six claudins were in loose contact with nCPE domains. For membrane embedment, the previously equilibrated membrane from state RC-VIIIa was used. Lipids that clashed with the newly added claudin subunits were either removed or refined. The system was generated, relaxed and simulated as before for 100 ns without constraints to obtain state RC-IXb. During the stepwise equilibration, the following residue clusters were constrained for an extended time (∼85 ns),

(i) Tyrosine cluster in cCPE – Y306, Y310 and Y 312,
(ii) Claudin residues fitting inside the pocket formed by the tyrosine cluster N149, P150, and L151,
(iii) Residues forming the linear-cis interface between claudins – L71, F147 and M160.

One well-equilibrated structure from the state RC-IXb model simulation was used to generate a variant where the comparatively tilted claudin monomers were straightened. The system was embedded in the membrane used for the RC-IXb model, lipids clashing with the claudins were removed or refined. The system was then relaxed, equilibrated with the aforementioned constraints, and finally simulated without constraints for 100 ns to obtain state RC-IXa. Coulombic interactions were calculated using the short-range cutoff method with a 9.0 Å cutoff. The RESPA algorithm was used to integrate the bonded interactions with a timestep of 2.0 fs.

### Analysis of the MD simulation trajectories

The MD simulation trajectories of the production runs were used for various quantitative analysis. Using the MD trajectory analysis tools in BioLuminate, key parameters such as residue-residue distances, and interactions (hydrogen bonds and salt bridges) were quantified. Solvent-Accessible Surface Area (SASA) data was extracted using Visual Molecular Dynamics^91^ (VMD, version 1.9.4a57, release 2022-04, https://www.ks.uiuc.edu/Research/vmd). The MDAnalysis tool (version 2.3.0)^92, 93^ was used to calculate the normalized contact time of the pore-lining residues with ions. Additionally, the HOLE program ^66, 94^ was used to analyze and visualize the pore. Protein backbone, considered for the RMSD analyses, includes the two carbons, nitrogen, oxygen and hydrogens of main chain of the residues.

### In vitro assays

#### CPE protein preparation

For the construction of a plasmid encoding CPE carrying a C’terminal StrepII tag (CPE-Strep) fusion protein, previously described pTrcHis-Topo-optCPEwt was used as a template.^15^With site-directed mutagenesis using a KLD enzyme mix the bases between 4285 and 4317, including the His-Tag, were removed and through Gibson Assembly a Strep-Tag was inserted at base 5313, at the C’terminal end of CPE. CPE was expressed similar as described previously^15^ with modifications mentioned in Supplementary Methods.

#### GST-cCPE protein preparation

GST-cCPE was expressed and purified as described previously.^36^ For details see Supplementary Methods.

### Cell culture

HEK293 (HEK) cells were cultured in DMEM-Earle’s media (Sigma-Aldrich, Germany) supplemented with 10% (v/v) fetal bovine serum, 100 U/ml penicillin and 100 μg/ml streptomycin (Sigma-Aldrich, Germany) at 37 °C in a humidified 5% CO2 atmosphere.

For Hek-CLDN3WT and Hek-CLDN3-S68E cells, previously described^68^, 0.5 mg/ml G418 was additionally added to the medium for selection.

### Cell viability assays

Cell viability assays were performed similar as described previously.^16, 64^ For details see Supplementary Methods.

### Cellular binding assays

Cellular binding assays were performed similar as described previously.^36^ For details see Supplementary Methods.

## Supporting information

Graphical Abstract

## Acknowledgments

This research was funded by the Deutsche Forschungsgemeinschaft (DFG, German Research Foundation) – DFG PI 837/4-3 – 289412825 and DFG GRK 2318 – 318905415 (A02) and the Sonnenfeld Stiftung, Berlin (hardware and licenses). We thank Alina Roderer for technical assistance.

## Author contributions

S.K.N.: Investigation, Formal analysis, Validation, Visualization, Methodology, Writing – review & editing; J.W.: Investigation, Formal analysis, Visualization, Methodology, Writing – review & editing; D.R.: Conceptualization, Supervision, Writing – review & editing; J.P.: Conceptualization, Supervision, Investigation, Visualization, Writing – original draft preparation, review & editing

## Notes

### Competing Interest Statement

The authors have declared no competing interest.

