## Supplementary figures and images for "C. perfringens enterotoxin-claudin pore complex: Models for structure, mechanism of pore assembly and cation permeability"

### Graphical Abstract

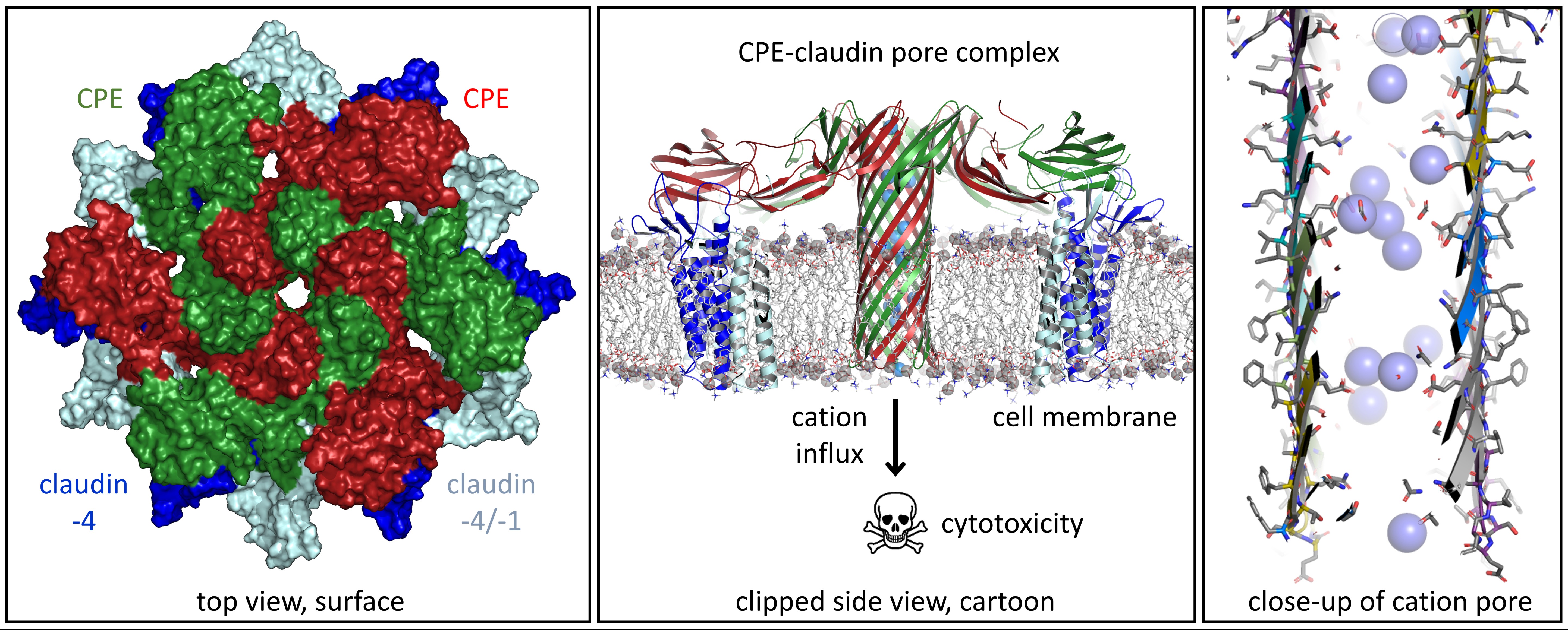
