## Supplementary Figures, Methods and Movie Legends for "C. perfringens enterotoxin-claudin pore complex: Models for structure, mechanism of pore assembly and cation permeability"

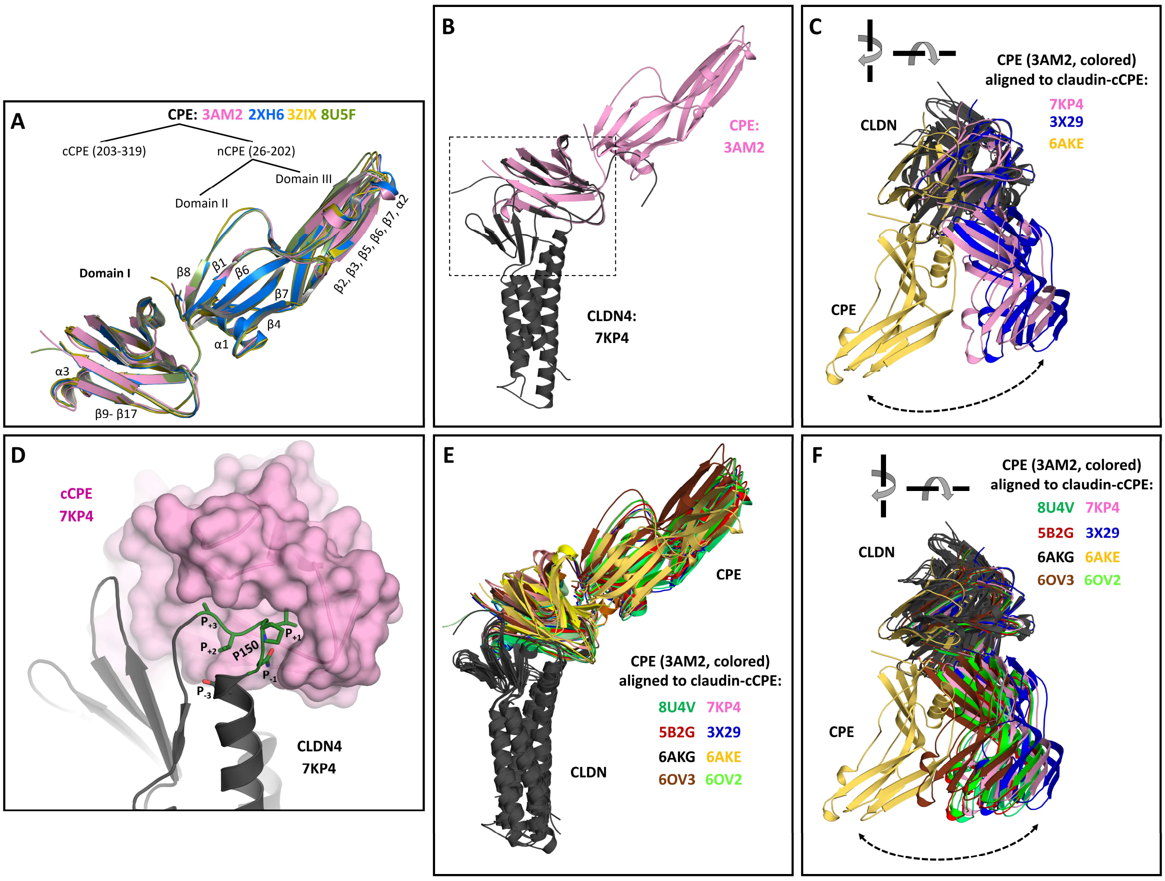


**Figure S1 (A)** Structural alignment of different CPE crystal structures. All monomer structures are very similar. Nomenclature of secondary structure elements and domains according to Kitadokoro et al.[1]. For further analysis 3AM2 was selected as a representative structure. **(B-F)** Structural alignment of CPE (3AM2) to different cCPE-claudin complex structures. Each claudin chains was aligned to the claudin-4 chain of 7KP4. Afterwards, the cCPE domain of a CPE chain was aligned to the respective cCPE chain of the cCPE-claudin complex. Compared to B/E the view in C/F is rotated by ~90° in two directions (arrows). **(B, E)** The different aligned CPE-claudin complexes differ not concerning the angle of CPE towards the membrane plane (approximately orthogonal to the axis of the transmembrane helix bundle of the claudin). **(C, F)** In contrast, the complexes differ with respect to the rotational angle around the axis of the transmembrane helix bundle of the claudin. **(C)** The cCPE-claudin complex structure used for the pore assembly model (7KP4) is shown in comparison to the two most extreme rotational variants (6AKE and 3X29). The comparison of the different CPE-claudin complexes suggests that the binding of the cCPE domain to claudins allows a certain degree of rotational flexibility even for CPE as a rigid body. **(D)** Close-up of dashed box in (B) with 7kp4 in slightly different perspective, highlighting the interaction between Cldn4-ECS2 and cCPE. The among classic claudins conserved P150 is used as a reference position. The positions $P_{-3}$, $P_{-1}$, P, $P_{+1}$, $P_{+2}$, $P_{+3}$ of the interaction motif ((D/E(P-4) x(P-3) x(P-2) N(P-1)P(P) L/M/V(P+1) V(P+2) P/A(P+3)) are indicated and the corresponding residues shown as sticks. The key region $P_{-1}$, to $P_{+3}$ is shown in green. cCPE is shown as transparent surface.

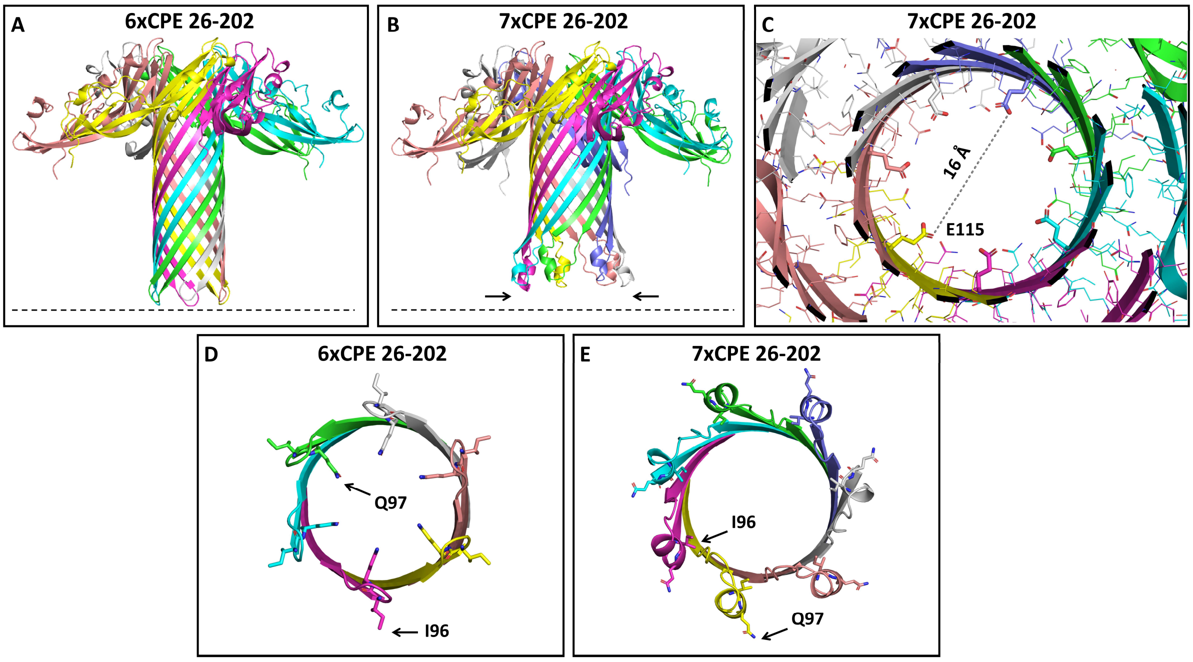


**Figure S2**: Comparison of Colabfold nCPE hexamer and nCPE heptamer models. (**A, B**) Side views of hexamer and heptamer. Note the inconsistent membrane-spanning region at the tip of the heptamer pore (arrows). (**C**) Constriction at E115 ring in the cap region of the β-barrel of the heptamer model (16 Å). E115 residues shown as sticks, others as lines. (**D, E**) Cross-sections through tips of the transmembrane barrel of the hexamer and heptamer. View from cytoplasmic side. I96 faces towards the hydrophobic membrane-embedded outside of the barrel in the hexamer and to the hydrophilic lumen in the heptamer. The opposite is observed for Q97. Lining, diameter and length of pore fits better to existing data for the hexamer.

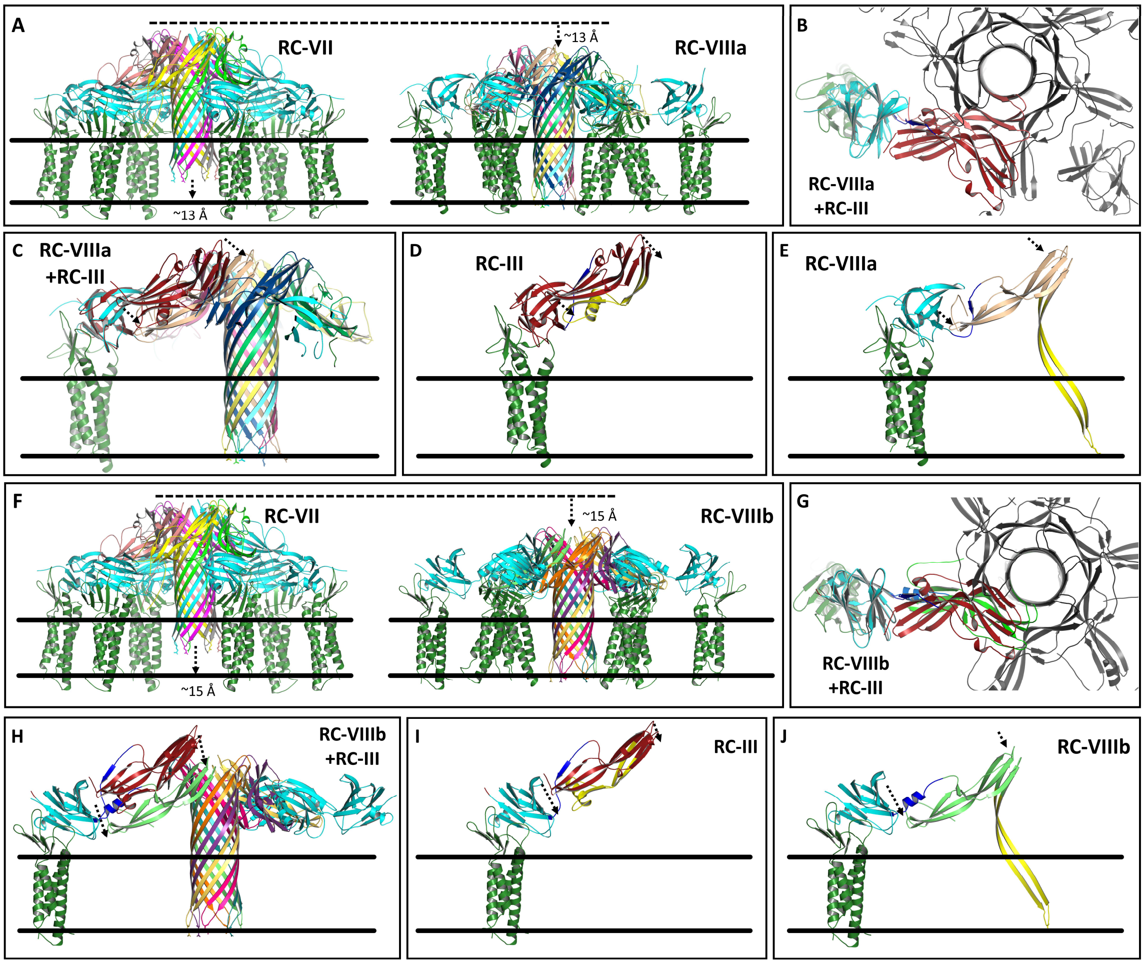


**Figure S3**: Conformation change in linker region between cCPE and nCPE (residues 191-205, blue) allowing full transmembrane penetration of β-hairpin tips of pore. Comparison of the two different conformational variants RC-VIIIa (**A-E**) and RC-VIIIb (**F-J**). (**A**) Comparison of CLDN4-bound CPE hexamer pore complexes without (state RC-VII) and with downwards shift of nCPE (state RC-VIIIa). E94 at β-hairpin tip is shown as stick. (**B**) Top view on RC-VIIIa with superimposed state RC-III to visualized the conformational difference between the monomeric CPE structure (3AM2 of RC-III, red) and CPE in RC-VIIIa (cyan, pink). (**C**) Clipped view of state RC-VIIIa focused on one claudin with superimposed state RC-III to visualized the conformational difference between the monomeric CPE structure (3AM2 of RC-III, red) and CPE in RC-VIIIA (cyan, beige). Individual CPE/claudin dimers are shown separately for RC-III (**D**) and RC-VIIIa (**E**). The region 73-116 that changes conformation is labeled in yellow in (D) and (E). In RC-III it includes the α1-helix, in RC-VIIIa it forms the main part of the pore β-barrel. The shift of nCPE in RC-VIIIa relative to the position in RC-III is labeled by dashed arrows in (C-E). **(F)** Comparison of CLDN4-bound CPE hexamer pore complexes without (RC-VII) and with variant downwards shift of nCPE (RC-VIIIb). (**G**) Top view on RC-VIIIb with superimposed state RC-III to visualized the conformational difference between the monomeric CPE structure (3AM2 of RC-III, red) and CPE in RC-VIIIb (cyan, green). (**H**) Clipped view of state RC-VIIIb focused on one claudin with superimposed state RC-III to visualized the conformational difference between the monomeric CPE structure (3AM2 of RC-III, red) and CPE in RC-VIIIb (cyan, green). Individual CPE/claudin dimers are shown separately for RC-III (**I**) and RC-VIIIa (**J**). The shift of nCPE in RC-VIIIb relative to the position in RC-III is labeled by dashed arrows in (H-J). Note that in RC-VIIIa part of the nCPE-cCPE linker region (blue) still forms a β-strand (E) similar as in 3AM2 of RC-III (D, I) whereas the corresponding linker region in RC-VIIIb forms an α1-helix above the β-sheet (β1, β6, β 7) in domain II (J).

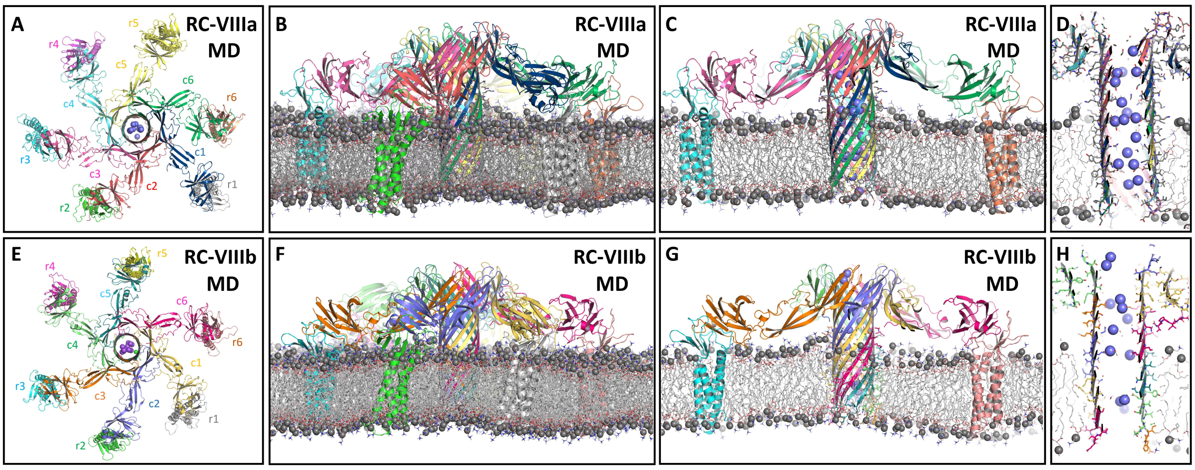


**Figure S4**: MD simulations of CPE-CLDN4 pore complex states RC-VIIIa (**A-D,** same images as in Figure 4) and RC-VIIIb (**E-H**). Comparison of the two variants. In both simulations the pore β-barrel outside and inside the membrane and the pore lining are well maintained. The pore β-barrel and the claudins are well embedded in the membrane and the cCPE-claudin interaction is stable. See also legend of Figure 4.

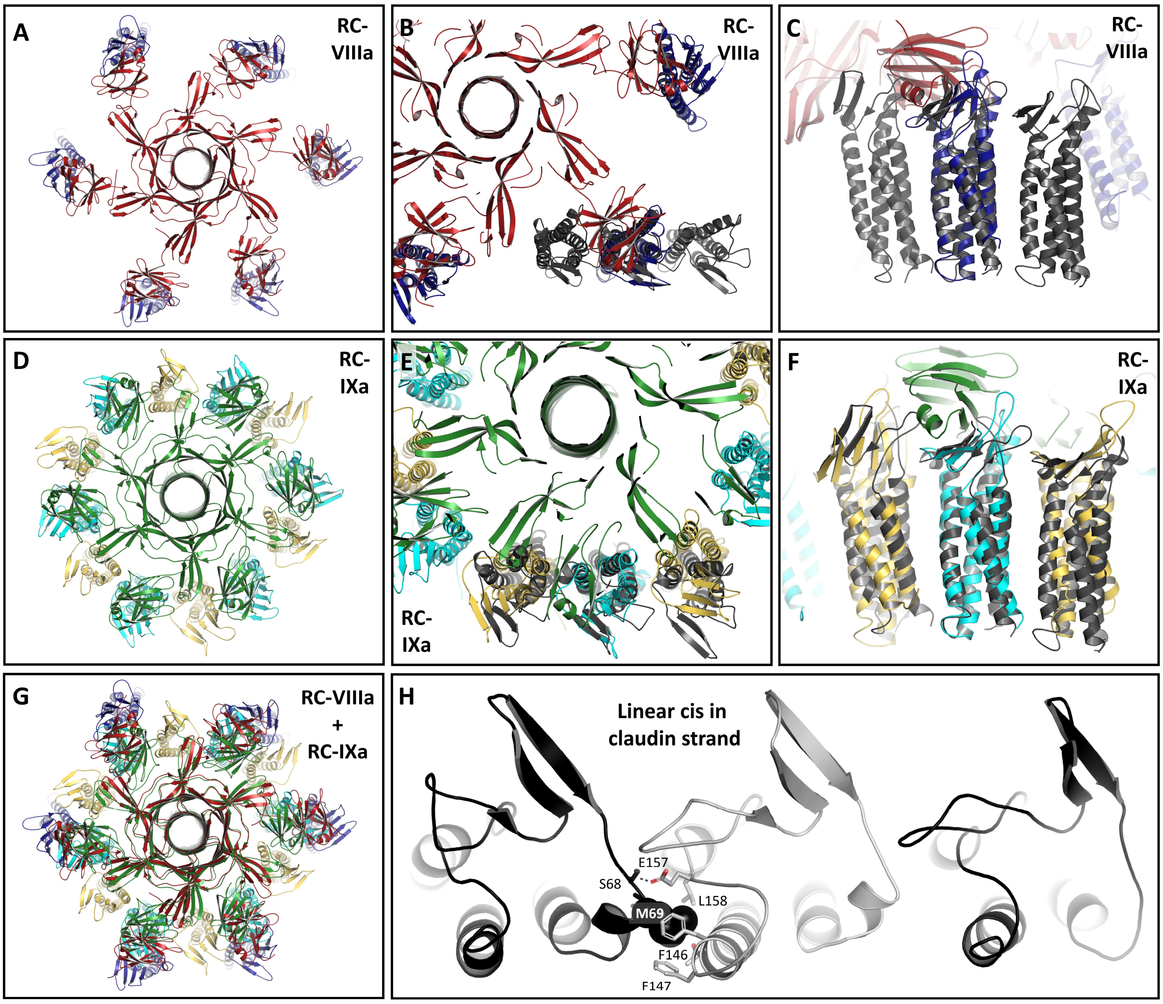


**Figure S5**: Manual addition of six more claudin subunits to generate a dodecameric claudin ring. (**A**) Top view of starting state RC-VIIIa with six CPE (red) and six claudin (blue) subunits. (**D**) Top view of resulting state RC-IXa with 6 CPE (green) and 12 claudin (cyan & yellow) subunits. (**G**) Top view of RC-VIIIa and RC-IXa superposition. The RC-IXa model was generated according to the following hypothesis: (1.) 12 claudins are part of the pore complex (biochemical evidence), and (2.) the interface between the claudins could be a variation of the linear-cis interface found in claudin strands. In (**H**) the linear-cis interface of a CLDN10b strand model[2] is shown. Conserved key interfacial residues F146 (corresponding to CLDN4-F147), F147 (corresponding to CLDN4-Y148), L158 (corresponding to CLDN4-M160), E157 (corresponding to CLDN4-E159), and S68 (corresponding to CLDN4-S69) are shown as sticks and M69 (corresponding to CLDN4-L71) as spheres. Conformational difference at cis-interfacial regions are caused by cCPE-binding (crystal structure evidence, [3-5]). In the model state RC-VIIIa the distance between CPE-bound claudins is too big to fit in a cis-interacting claudin (**B**), top view and (**C**), side view of RC-VIIIa with superimposed linear cis-claudin trimer (gray). For fitting, a planar CLDN4 dodecameric ring was manually docked based on 30° rotations of the linear-cis interface, (**D**) top view, (**E**) top view and (**F**) side view with superimposed linear cis-claudin trimer (gray). Six cCPE subunits were docked to each second CLDN4 subunit according to cCPE/CLDN4 structure 7kp4. nCPE was positioned close to free CLDN4 subunits. This resulted in a shifted and turned position of the cCPE domain relative to the nCPE domain (see Figure 5). Both domains were connected by manually modifying the nCPE-cCPE linker region including the β8-strand).

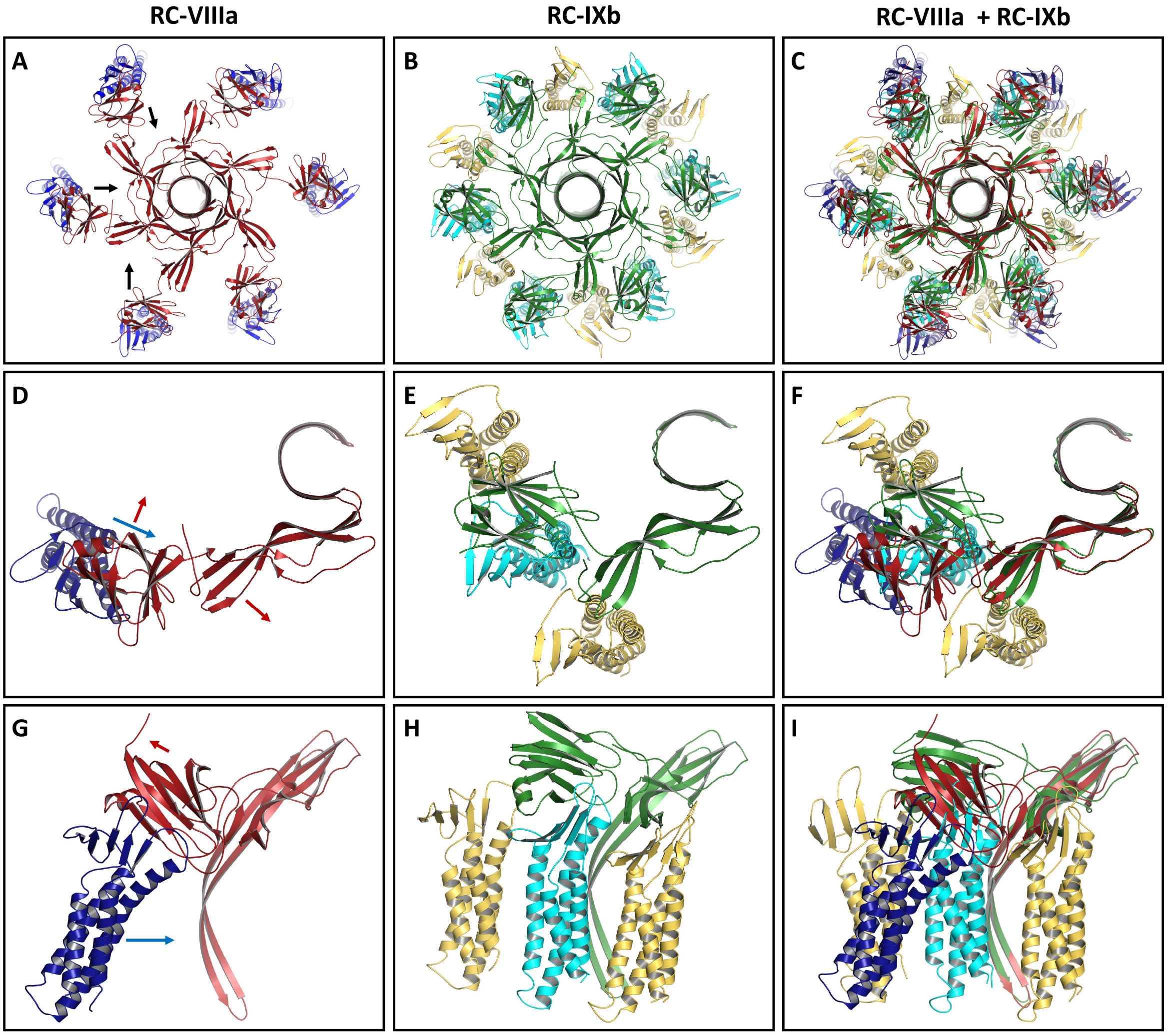


**Figure S6**: Details of modification of relative cCPE/nCPE positions for state RC-IXb. (**A-C**) Top views. (**A**) State RC-VIIIa: CPE pore hexamer (red) bound to six individual claudins (blue). (**B**) State RC-IXb: CPE pore hexamer (green) bound to a dodecameric claudin ring. Claudin subunits primarily bound to CPE are colored cyan, additional claudin subunits are colored yellow. (**C**) Superposition of RC-VIIIa and RC-IXb. (**D-I**) Single CPE subunits bound to claudins are shown in top view (**D**, RC-VIIIa), (**E**, RC-IXb), (**F**, both) or side view (**G**, RC-VIIIa), (**H**, RC-IXb), (**I**, both). For RC-VIIIa, one CPE subunit bound one claudin subunit. For RC-IXb, in addition to the primarily CPE-bound claudin subunit, both adjacent claudins are shown. The arrows indicate direction of movement of cCPE (red) and claudin (blue) from state RC-VIIIa to state RC-IXb. 100 ns snapshots of MD simulation are shown.

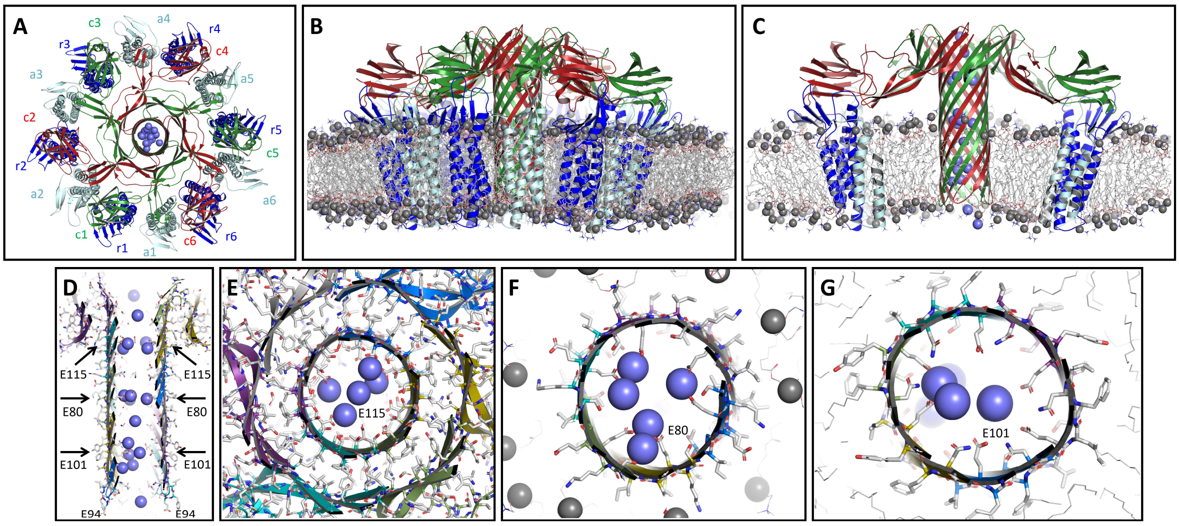


**Figure S7**: MD simulation of CPE hexamer anchored to dodecameric CLDN4 ring (pore complex state RC-IXb). Shown are snapshots after 100 ns of free simulation with protein as cartoon, relevant residues as sticks, lipids acyl chains as gray lines and phosphate head groups as gray spheres, sodium ions as blue spheres. **(A)** Top view with CPE (c1-c6, green & red), canonical receptor CLDN4 (r1-r6, blue) and additional CLDN4 (a1-a6, cyan) subunits. (**B, C**) Side view (clipped in (C)). The pore barrel and the pore cap are well preserved. The pore complex is well embedded in the membrane. (**D**) Clipped side view of pore to illustrate pore lining and strong presence of sodium ions in the pore lumen. (**E**) The inner and outer β-barrel in the cap region (upper arrow in (D)) are held together mainly by interactions between hydrophobic residues. In the center, ring of six E115 residues strongly attracts cations (spheres). Rings formed by six E80 residues (**F**) slightly above membrane plane (middle arrow in (D)) and by six E101 residues (lower arrow in (D)) within the membrane plane (**G**) also strongly attract cations.

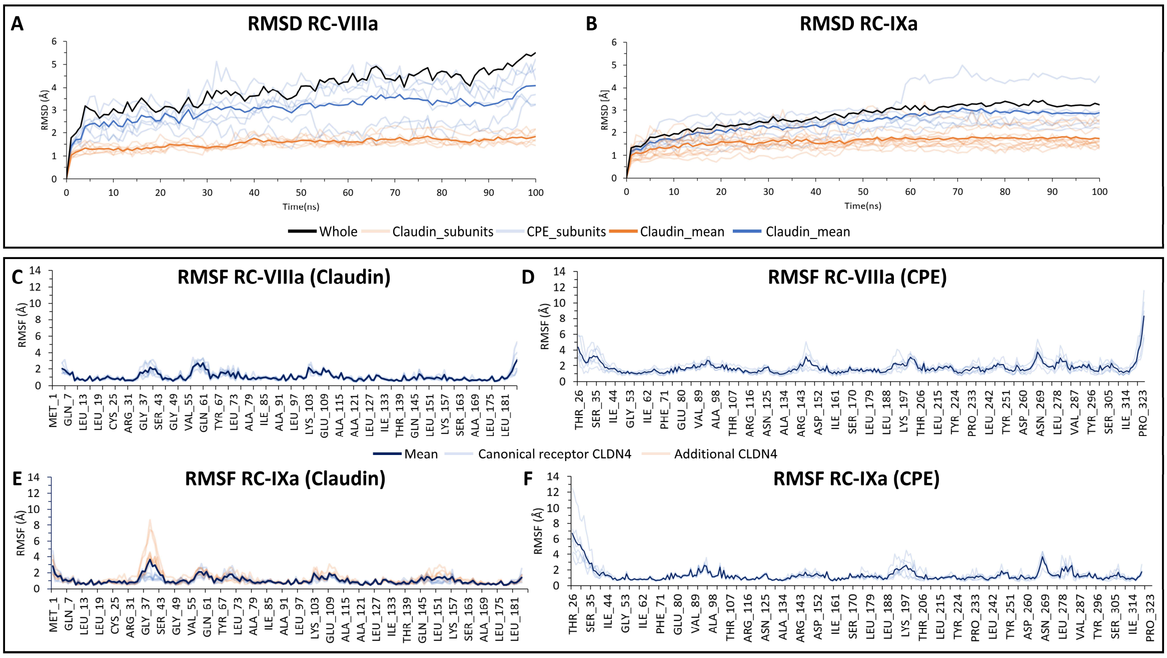


**Figure S8: (A, B)** Root-mean-square deviation (RMSD) of protein backbone for 6-claudin comprising RC-VIIIa and 12-claudin comprising RC-IXa pore complex states respectively. The change in RMSD over the simulation time (100 ns) with respect to the starting structure (0 ns) is plotted. RMSD of the backbone of the whole RC-VIIIa (including both CPE and claudins, shown in black) was varying slightly more than of the RC-IXa. For RC-VIIIa, the discontinuous six claudin ring resulted in higher flexibility as the RMSD varied until ~5.5 Å. On the contrary, for RC-IXa with the presence of 12-claudin ring, the RMSD reached saturation (at ~3.0 Å) after ~50 ns. Notably, the backbone of claudins (orange lines) in both complexes deviated around ~1.5 Å, which is comparable to the RMSD values of claudin backbones in previously published claudin strand models.[2, 6] Hence, the mean RMSD of the backbone of CPE subunits (blue lines) which fluctuated around ~3.0 Å and ~2.4 Å, for RC-VIIIa and RC-IXa respectively, resulted in the variation in the whole RMSD between both complexes. **(C** to **F)** Mean root mean square fluctuation (RMSF) of the amino acid residues of claudins and CPEs in both complexes are given. In both, RC-VIIIa and RC-IXa (**C** and **E**), the fluctuation of residues in the claudins were mostly similar, but differing mainly in the β1-β2 loop (residues 36-42) and ECS2 (residues 146-156) regions. To understand the difference, the RMSF values for canonical receptor CLDN4 and additional CLDN4 subunits in RC-IXa are given separately in addition (**E**). Here, we could observe the noticeable fluctuation in the residues of these regions for the additional CLDN4 subunits. Similarly, in the RMSF plots of CPE subunits and their mean (**D** and **F**), less deviations could be seen in certain regions in CPE of RC-IXa (residues 142 to 152), mainly due to the loose contacts of CPE subunits with the additional CLDN4 subunits, which were not present in RC-VIIIa as there were no claudins in the gap. Expectedly, the manually-modified nCPE-cCPE linker segment (V187-A204) in RC-IXa fluctuated slightly more than the average. A similar noticeable higher fluctuation could be seen in the segment around N269, since it did not take part in any key intermolecular or intramolecular interaction. Overall, the mean RMSF for most residues in both claudins and CPEs were < 1.5 Å which indicates their stability.

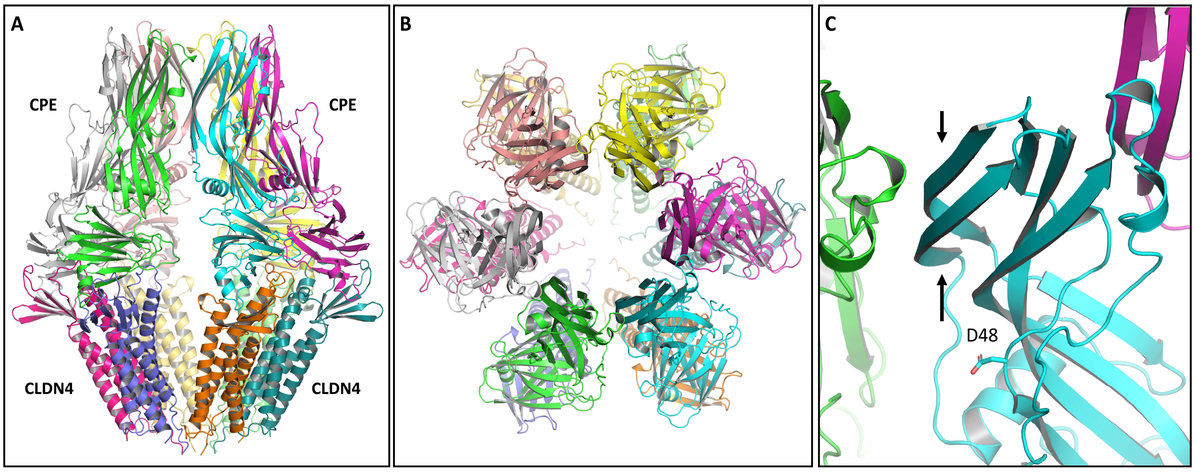


**Figure S9:** Example Alphafold3 output. CPE and claudin subunits are shown as colored cartoons. **(A)** Side view. **(B)** Top view. **(C)** Close up. β3- and β5-strands (arrows) in nCPE are not forming a β-barrel, in contrast to all Alphafold2-based CPE states reported in this study. D48 is not participating in an interface between neighboring CPE subunits. Six CPE-(26-319) and six hCldn4 (1-191) molecules were given as protein inputs using the AlphaFold3 Server (https://golgi.sandbox.google.com/).

**Movie S1**: Conformational changes of nCPE hexamer model. This movie shows a morphing from the pre-pore state nCPE-IV to pre-pore state nCPE-V. View from extracellular side. CPE subunits shown as cartoon in different color. See Figure 1 for more details.

**Movie S2**: Conformational changes of nCPE hexamer model. This movie shows a morphing from the pre-pore state nCPE-V to pore state nCPE-VII. View from extracellular side. CPE subunits shown as cartoon in different color. See Figure 1 for more details.

**Movie S3**: Conformational changes of nCPE hexamer model. This movie shows a morphing from the pre-pore state nCPE-IV to pore state nCPE-VII. View from extracellular side. CPE subunits shown as cartoon in different color. See Figure 1 for more details.

**Movie S4**: MD simulation for pore state nCPE-VII. Last 50 ns of simulation are shown. CPE subunits shown as cartoon in different colors, $Na^+$ and $Cl^-$ ions are shown as red and green spheres, respectively. Membrane and water are not shown for clarity. Side view (large) and top view from extracellular side (small). The hexameric β-barrel pore is stable throughout the simulation and strongly attracts $Na^+$ ions.

**Movie S5**: MD simulation for pore state RC-VIIIa. Last 50 ns of simulation are shown. CPE and claudins subunits are shown as cartoon in different colors, $Na^+$ and $Cl^-$ ions are shown as red and green spheres, respectively. Membrane lipids are shown as lines with phosphate atoms as pink spheres. Clipped side view (large) and full top view from extracellular side (small). The complex of six CPE and six CLDN4 subunits including the hexameric CPE β-barrel pore is stable throughout the simulation. The pore strongly attracts $Na^+$ ions.

**Movie S6**: MD simulation for pore state RC-IXa. Last 50 ns of simulation are shown. CPE and claudins subunits are shown as cartoon in different colors, $Na^+$ and $Cl^-$ ions are shown as red and green spheres, respectively. Membrane lipids are shown as lines with phosphate atoms as pink spheres. Clipped side view (large) and full top view from extracellular side (small). The complex of six CPE and 12 CLDN4 subunits including the hexameric CPE β-barrel pore is stable throughout the simulation. The pore strongly attracts $Na^+$ ions.

**Supplementary Methods**

**CPE protein preparation**

CPE-Strep was expressed in *Escherichia coli* Rosetta-2. After expression was induced using 0.5 mM IPTG and subsequently 4 h of incubation at 37°C, CPE was purified from lysates using Strep-Tactin®XT columns (IBA, Göttingen, Germany). Bacteria from 1 l culture volume were harvested by centrifugation (12 min; 4500xg; 4°C) and resuspended in 50 ml lysis buffer (100 mM Tris/HCl; 150 mM NaCl; 1 mM EDTA, pH 8.0) additionally containing protease inhibitor cocktail (Merck). Lysis was performed with a LM10 Microfluidizer (Microfluidics, Westwood, USA) on ice. Cell debris was removed by centrifugation (30 min; 20000×g; 4°C) and supernatant loaded onto columns containing 2 mL Strep-Tactin® Superflow® high capacity resin (IBA, Göttingen, Germany). The column was washed (with 10 ml lysis buffer) and CPE eluted with lysis buffer containing additional 50 mM Biotin. The protein concentration was determined by NanoDrop Microvolume UV-Vis Spectrophotometers based on Protein A280.

**GST-cCPE protein preparation**

Plasmids carrying the open reading frame for GST-cCPEwt194−319 (GST-cCPE) have been previously documented.[7] Fusion proteins of GST-cCPE from these plasmids were produced in E. coli BL21 and purified according to established methods. The bacteria were cultivated until they reached an optical density of 0.6–0.8, at which point protein expression was triggered by adding 1 mM isopropyl-β-D-thiogalactopyranoside. Three hours after induction, the bacterial cells were collected and lysed in lysis buffer consisting of phosphate-buffered saline (PBS) with 1% (v/v) Triton X-100, 0.1 mM phenylmethylsulfonyl fluoride, and 1 mM ethylenediaminetetraacetic acid, along with a protease inhibitor cocktail (Merck). The cells were then sonicated with 15 pulses of 1 second each using a Vibra Cell Model 72434 BioBlock Scientific sonicator. The insoluble debris was separated by centrifugation at 20,000 × g for 30 minutes at 4°C. The GST-proteins were purified from the supernatant using glutathione agarose (Sigma-Aldrich) and subsequently dialyzed against PBS. The protein concentration was measured using the Pierce™ BCA Assay Kit (Thermo Fisher Scientific, Waltham, Massachusetts, USA).

**Cell viability assays**

To analyze CPE-mediated cell damage on Hek293 cells expressing the CLDN3-S68E mutant in comparison to Hek293-CLDN3WT cells, a viability assay was performed using 4,5-Dimethylthiazol-2-yl)-2,5-diphenyltetrazolium bromide (MTT). Per Well 5*10^4 cells were seeded on 96-well plates coated with PLL and kept at 37°C, in 5% CO2. At about 95 % confluence (after 24 h) the cells were incubated for 1 hour with variable dilutions of CPE or 0.01% (v/v) Triton X-100 (neg. control) before subsequently medium was replaced by medium without phenol red containing MTT (1.25 mM). After an additional 3-hour incubation, extraction and solubilization of water-insoluble blue-violet formazan was performed by treatment with 5% (v/v) Triton X-100 in 2-propanol for 20 minutes. Absorption was measured at 560 nm. The sample size was set to 6. For each concentration of each assay three technical replicates (n=3) were prepared. The half-maximal effective concentration (EC50) values were calculated from the normalized data with the GraphPad Prism software using the model "log(agonist) vs. response" model (four parameter). A paired *t*-test was performed to determine significant differences between the EC50 values of Hek293-CLDN3-WT and Hek293-CLDN3-S68E.

**Cellular binding assays**

3*10^5 cells were seeded to PLL-coated 24-well plates and after 24 hours of cultivation at 37°C, in 5% CO2, the assay was performed at a confluence of about 90 to 100 %. Medium was exchanged with variable dilutions of cCPE (0, 2, 4, 8, 16, 32, 64, 128, 256, 512 nM) in cell culture medium (0.5 ml per well). After a 30 min incubation at 37 °C, in 5 % $CO_2$, cCPE was removed and the cells were fixed (4% [w/v] paraformaldehyde, 10 min), followed by washing with PBS. For the subsequent quenching, 0.25 ml quenching buffer (0.1 M Glycin in PBS) was added to each well. Bound GST-cCPE was detected via PhycoLink® anti-GST-R-phycoerythrin conjugate; or alternatively in one case with anti-GST first antibody and Alexa594-linked 2$^{nd}$ antibody. The signal was normalized to cell number (Hoechst 33342). Each well was treated with 0.25 ml blocking reagent (1%(w/v) BSA, 0.05 %(v/v) Tween-20 in PBS) with PhycoLink® anti-GST-R-phycoerythrin conjugate (1:250) as well as Hoechst 33342 (2 µM) and incubated for 1h; or alternatively with 0.25 ml blocking reagent with the first antibody (1:250) and subsequently after 1h incubation, with 0.25 ml blocking reagent with the secondary antibody (1:500) and Hoechst 33342 (2 µM), followed by another 1h incubation. Normalized fluorescence intensity of bound anti-GST antibody was plotted against GST-cCPE concentration for CLDN3WT as well as the mutant, CLDN3-S68E.The $K_D$ was calculated using nonlinear regression analysis for a single-site, specific binding in GraphPad Prism version 7.0 (San Diego, CA, USA). Unspecific binding was accounted for by subtracting the fluorescence signal after incubation of untransfected HEK293 cells with respective concentrations of GST-cCPE. Finally, a *t*-test was performed to determine significant differences between $K_D$ values of GST-cCPE binding to Hek293-CLDN3WT or Hek293-CLDN3-S68E. For each assay two technical replicates of each well were prepared, and the assay was performed three times.
